# Epithelial HVEM promotes basement membrane synthesis and intraepithelial T cell survival and migration

**DOI:** 10.1101/2020.09.30.317628

**Authors:** Daisuke Takahashi, Qingyang Wang, Goo-Young Seo, Jr-wen Shui, Zbigniew Mikulski, Paola Marcovecchio, Masumi Takahashi, Charles D. Surh, Hilde Cheroutre, Mitchell Kronenberg

**Affiliations:** La Jolla Institute for Immunology, La Jolla, CA 92037, USA; Division of Biochemistry, Department of Pharmaceutical Sciences, Faculty of Pharmacy, Keio University, Tokyo 105-0011 Japan; Beijing Institute of Brain Sciences, Beijing 100850, P. R. China; Academic Road, Nankang, Institute of Biomedical Sciences, Academia Sinica, Taipei 11529, Taiwan; Institute for Basic Science (IBS), Academy of Immunology and Microbiology, Pohang, South Korea; Division of Biology, University of California San Diego, La Jolla, CA 92037, USA

**Author notes:** co-First author (Daisuke Takahashi, Qingyang Wang, Goo-Young Seo).

**Keywords:** HVEM, intraepithelial T cells, intestinal epithelial cells, basement membrane, cell survival

## Abstract

Intraepithelial T cells (IET) provide continuous surveillance of the intestinal epithelium, but little was known about how epithelial-derived signals regulate the IET population. We show that epithelial expression of the herpes virus entry mediator (HVEM), a member of the TNF receptor superfamily (TNFRSF), maintained the survival of small intestine IET, especially innate-like TCRαβ^+^ cells lacking CD4 and CD8β. Patrolling movement of all CD8α^+^ IET also was impaired in the absence of HVEM. HVEM-deficient epithelial cells exhibited downregulation of synthesis of basement membrane components, including collagen IV. Collagen IV supported IET survival *in vitro* via interactions with β_1_ integrins expressed by the IET; absence of β_1_ integrins decreased some IET subsets. Therefore, these data define a circuit whereby epithelial cells regulate intestine resident T lymphocyte populations through basement membrane synthesis.

## Introduction

Intestinal intraepithelial lymphocytes (IEL) are one of the largest population of lymphocytes in the body. They are found above the basement membrane within the intestinal epithelium, and they interact extensively with intestinal epithelial cells (IEC) by actively patrolling the basement membrane and by migration into the lateral intercellular space (Edelblum et al., 2012; Hoytema van Konijnenburg et al., 2017; Sumida et al., 2017; Wang et al., 2014). IEL are believed to be crucial for maintenance of integrity of the intestinal barrier, repair of wounds and protection from the pathogenic invasion (Cheroutre et al., 2011). They include ILC but are mostly T lymphocytes and we refer to those T cells here as intraepithelial T cells (IET). On the basis of development and antigen-recognition, distinct IET subsets have been divided into two groups termed “induced” IET, or type a and “natural” IET or type b (Cheroutre et al., 2011; Hayday et al., 2001; Olivares-Villagomez and Van Kaer, 2018).

Induced IET express αβ TCRs together with one of the TCR co-receptors that promote TCR signaling, either CD4 or CD8αβ. They are tissue resident memory cells derived from antigen-primed conventional or mainstream CD4^+^ or CD8αβ^+^ T cells (Lefrancois and Masopust, 2002; Masopust et al., 2001), but have an oligo clonal TCR repertoire (Regnault et al., 1994). By contrast, natural or type b IET express either αβ TCRs or γδ TCRs, but lack expression of the TCR co-receptors, CD4 and CD8αβ. Natural IET are approximately 50% of the IET population, and they can be either CD4 and CD8α double negative (DN), but they often express CD8αα homodimers, which lack TCR co-receptor function (Cheroutre and Lambolez, 2008). Natural IET TCRαβ^+^ precursors undergo an alternative, self-antigen-based or agonist thymic selection and maturation process, followed by direct migration to the intestinal epithelium (Leishman et al., 2002; Oh-Hora et al., 2013; Yamagata et al., 2004). Moreover, natural TCRαβ^+^ IET have a distinct set of specificities compared to TCRαβ^+^ CD8αβ lymphocytes (Mayans et al., 2014; McDonald et al., 2014; McDonald et al., 2018). The functions of natural IET have not been fully characterized, however, TCRγδ^+^ IET have been implicated in fast clearance and repair of damaged epithelium by various mechanisms (Boismenu and Havran, 1994; Guy-Grand et al., 1998; Komano et al., 1995) and there are reports that both natural IET populations respond to infections (Hoytema van Konijnenburg et al., 2017; Ismail et al., 2011; Saurer et al., 2004). Moreover, the factors important for the maintenance of natural IET in the epithelium have not been completely defined, but the aryl hydrocarbon receptor (AHR) (Li et al., 2011), T-bet (Klose et al., 2014) and MyD88 (Yu et al., 2006) have been implicated for both populations, microbial sensing is important for TCRαβ^+^ natural IET (Bandeira et al., 1990; Hoytema van Konijnenburg et al., 2017; Umesaki et al., 1993) and G protein-coupled receptor 18 (GPR18) positively regulates TCRγδ^+^ IET (Wang et al., 2014) while GPR55 is a negative regulator of these cells (Sumida et al., 2017). IL-15 contributes to natural IET maintenance, especially TCRγδ^+^ IET, providing an example of an epithelial cell-derived influence (Hu et al., 2018; Lodolce et al., 1998; Ma et al., 2009).

HVEM is a member of the tumor necrosis factor receptor superfamily (TNFRSF14) expressed by intestinal epithelial cells, lymphocytes and other cell types. It has been shown to be important for the regulation of barrier or mucosal immunity in mice in the context of infection or inflammation in a variety of contexts (Breloer et al., 2015; Desai et al., 2018; Flynn et al., 2013; Herro et al., 2018; Seo et al., 2018; Shui et al., 2012; Steinberg et al., 2013; Steinberg et al., 2008). Although HVEM is expressed constitutively by intestinal epithelial cells, a function for HVEM at steady state in the mucosae had not been described. Here we show that interactions of HVEM expressed by small intestine (SI) epithelial cells are important for the homeostasis and patrolling function of IET subsets, particularly for natural IET. We identify a mechanism for this unique lymphoid-epithelial interaction, which includes HVEM-stimulated synthesis of extracellular membrane components that affect natural IET in part through extracellular membrane interactions with integrins.

## Results

### HVEM is important for maintaining IET

To determine if HVEM functions in the regulation of IET in the intestine at steady state, we used flow cytometry to examine the total CD45^+^ cell number from mice with a germline deletion of the *Hvem* (*Tnfrsf14*) gene. Total CD45^+^ cells were significantly reduced throughout the SI, but not in the cecum and colon (Figure 1A). We also observed a decrease in CD45^+^ cells throughout the SI epithelium in *Hvem*^-/-^ mice with confocal microscopy (Figures 1B, 1C). Most of the CD45^+^ cells in the epithelium are T lymphocytes, including TCRγδ^+^ cells and TCRαβ^+^ cells, with a smaller population of ILC1 cells (Fuchs et al., 2013). We gated on the different IET subsets as shown in Figure S1A. Considering the different subsets, TCRαβ^+^CD8αα^+^ IET exhibited the most pronounced reduction in cell frequency (Figure 1D) and cell number (Figure 1E) in the proximal SI, but also elsewhere in the SI from *Hvem*^-/-^ mice (data not shown). TCRγδ^+^CD8αα^+^ were also affected in the proximal and middle portions of the SI. There was also some decrease in induced IET, for example decreased TCRαβ^+^CD8αβ^+^ IET in the proximal SI (Figure 1E). These data indicate that HVEM is important for the accumulation of IET at steady state, particularly TCRαβ^+^CD8αα^+^ IET.

**Figure 1.**
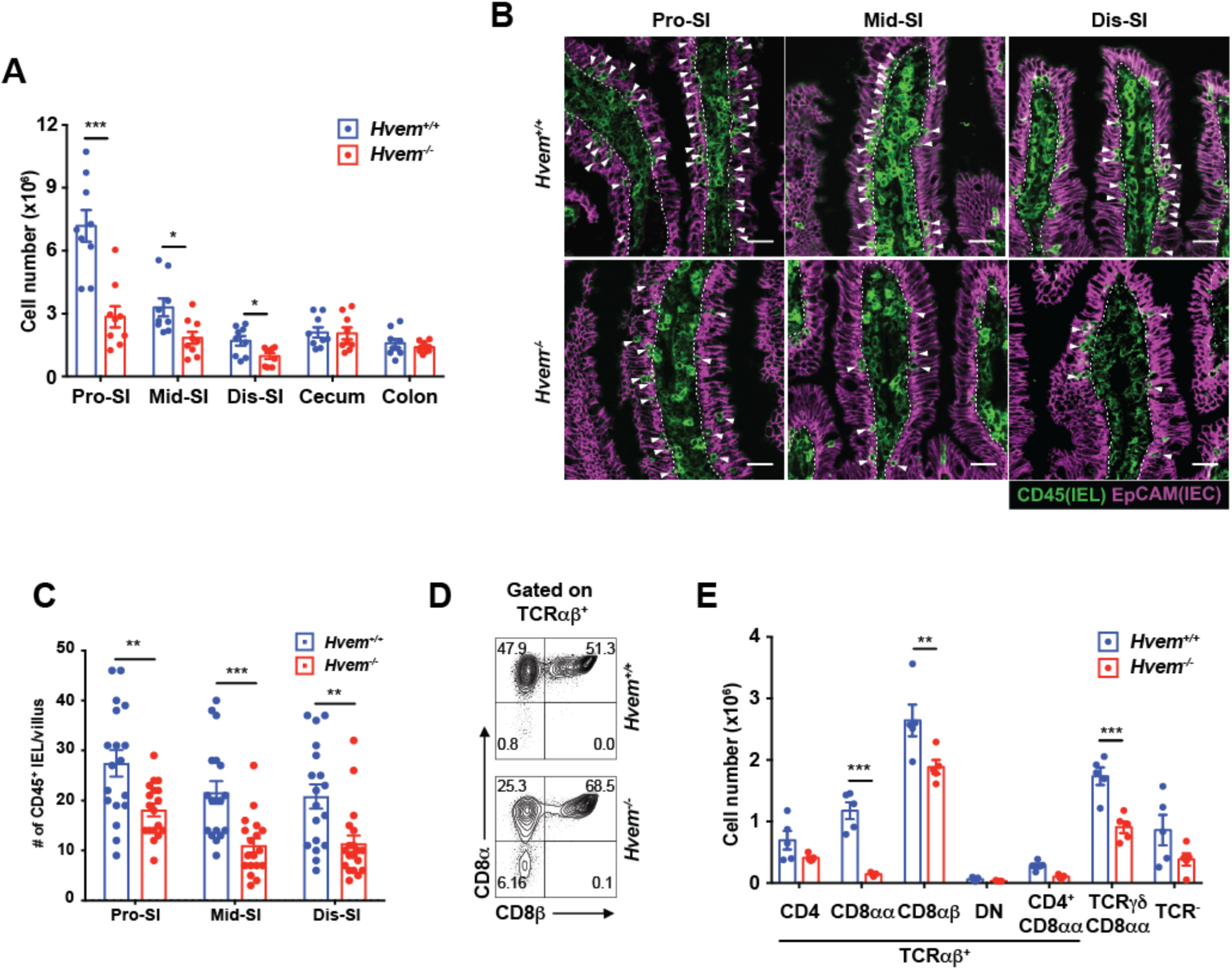
HVEM is important for maintaining IET. (A) Total IEL numbers by flow cytometry in proximal SI (Pro-SI), middle SI (Mid-SI), distal SI (Dis-SI), cecum and colon from *Hvem*^*+/+*^ and *Hvem*^-/-^ mice. (B) Representative immunofluorescence staining of CD45^+^ cells from the SI in *Hvem*^*+/+*^ and *Hvem*^-/-^ mice. White arrowheads indicate CD45^+^ intraepithelial cells (IEL) in the epithelium. Dashed white lines indicate the interface between the epithelium and lamina propria. Scale bars, 25μm. (C) Quantification of CD45^+^ IEL in villi from *Hvem*^*+/+*^ and *Hvem*^-/-^ mice. (D) Representative plots of TCRαβ^+^CD8αα^+^ and TCRαβ^+^CD8αβ^+^ IET in TCRαβ^+^ IET from proximal SI in *Hvem*^*+/+*^ and *Hvem*^-/-^ mice. (E) Absolute numbers of the indicated subsets in total IEL from proximal SI in *Hvem*^*+/+*^ and *Hvem*^-/-^ mice. Statistical analysis was performed using an unpaired t-test (A, C, E). Statistical significance is indicated by *, p < 0.05; **, p < 0.01; ***, p < 0.001. In A and E, bars show the mean and each symbol represents a measurement from a single mouse. In C, bars show the mean and each symbol represents cell numbers of CD45^+^ IEL per a boundary approximating a villus. Data represent pooled results from at least two independent experiments with at least four mice per group in each experiment (A), compiled from four independent experiments (B, C) or representative results from one of at least two independent experiments with at least four mice in each experimental group (E). Groups of co-housed mice were analyzed. See also Figure S1.

### Epithelial HVEM is required for CD8αα^+^ IET

Previous work identified the importance of HVEM expression by different cell types in mucosal immunity during infections and colitis (Schaer et al., 2011; Seo et al., 2018; Shui et al., 2012; Steinberg et al., 2008; Wang et al., 2005). Considering the importance of HVEM for TCRαβ^+^ CD8αβ^+^ memory and resident memory T cell formation (Flynn et al., 2013; Steinberg et al., 2013), we tested for a T cell intrinsic effect of HVEM expression. We analyzed the number of double-positive TCRαβ^+^ (DN TCRαβ^+^) thymocytes, which are precursors of CD8αα^+^TCRαβ^+^ IET (Cheroutre et al., 2011; Gangadharan et al., 2006) and found no difference between wild type and *Hvem*^*-/-*^ mice (Figure S1B). We transferred these precursor cells, NK1.1^-^TCRγδ^-^TCRαβ^+^ thymocytes (Gangadharan et al., 2006), into *Rag1*^*-/-*^ mice. A 1:1 co-transfer of donor thymocytes from *Hvem*^-/-^ mice (CD45.2) and congenic, wildtype CD45.1^+^ C57BL/6 mice was performed (Figure S1C). There was no defect in the ability of the *Hvem*^-/-^ donor thymocytes to give rise to TCRαβ^+^CD8αα^+^ IET. In fact, the *Hvem*^-/-^ donor cells were more highly represented at four weeks after transfer but the populations were equivalent at six weeks (Figure S1C). We also analyzed mice with a T cell-specific deletion of the *Hvem* gene driven by CD4-Cre. These *Cd4*-cre × *Hvem*^*fl/fl*^ (*Hvem*^*ΔCD4*^) mice showed no difference in the numbers of most IET subsets in the proximal SI (Figure S1D), although the CD4^+^CD8αα^+^ IET that are part of the CD4^+^ cytotoxic T lymphocytes (CD4^+^ CTL) population (Mucida et al., 2013) were reduced. Considering these results, a role for T cell HVEM expression is not the major influence in determining the size of the IET populations.

To test for the importance of HVEM at steady-state in another cell type, we examined HVEM expression by IEC. HVEM is predominantly on the basolateral surface of epithelial cells (Figure S2A), an optimal position for contact with immune cells, and it is expressed at a higher level in IEC from the proximal SI than the middle portion (Figure S2B). Interestingly, HVEM expression by IEC was not increased during infection (Shui et al., 2012) nor was it diminished in IEC from germ-free mice (Figure S2C). Also, HVEM was expressed in proximal small intestinal organoid cultures (Figure S2D, S2E). Therefore, constitutive HVEM expression by IEC was unaffected by acute infection or the absence of either the microbiota or IEL.

To test if there is a role for epithelial cell-expressed HVEM, we crossed *Villin*-cre mice with *Hvem*^*fl/fl*^ mice to generate mice with HVEM expression ablated only in epithelial cells (*Hvem*^*ΔIEC*^ mice, Figure S2F, S2G). Flow cytometric analyses indicated that *Hvem*^*ΔIEC*^ mice had significantly decreased CD45^+^ cell numbers in the proximal and middle small intestine compared to *Hvem*^*fl/fl*^ controls (Figure 2A). When the proximal SI was analyzed in more detail, the TCRαβ^+^CD8αα^+^ and TCRγδ^+^CD8αα^+^ IET subsets were decreased (Figures 2B, 2C), similar to *Hvem*^*-/-*^ mice. TCRαβ^+^CD8αα^+^ IET were also decreased in the middle and distal SI (Figure S2H) and TCRγδ^+^CD8αα^+^ IET were decreased in the middle portion (Figure S2H). The CD4 CTL marked by CD8α expression were decreased in parts of the SI as well, suggesting there could be HVEM-dependent contributions to this population that are both T cell-intrinsic and epithelial cell dependent. These effects were specific to the SI, however, as no IET populations were diminished in the colon of *Hvem*^*ΔIEC*^ mice (Figure S2H).

**Figure 2.**
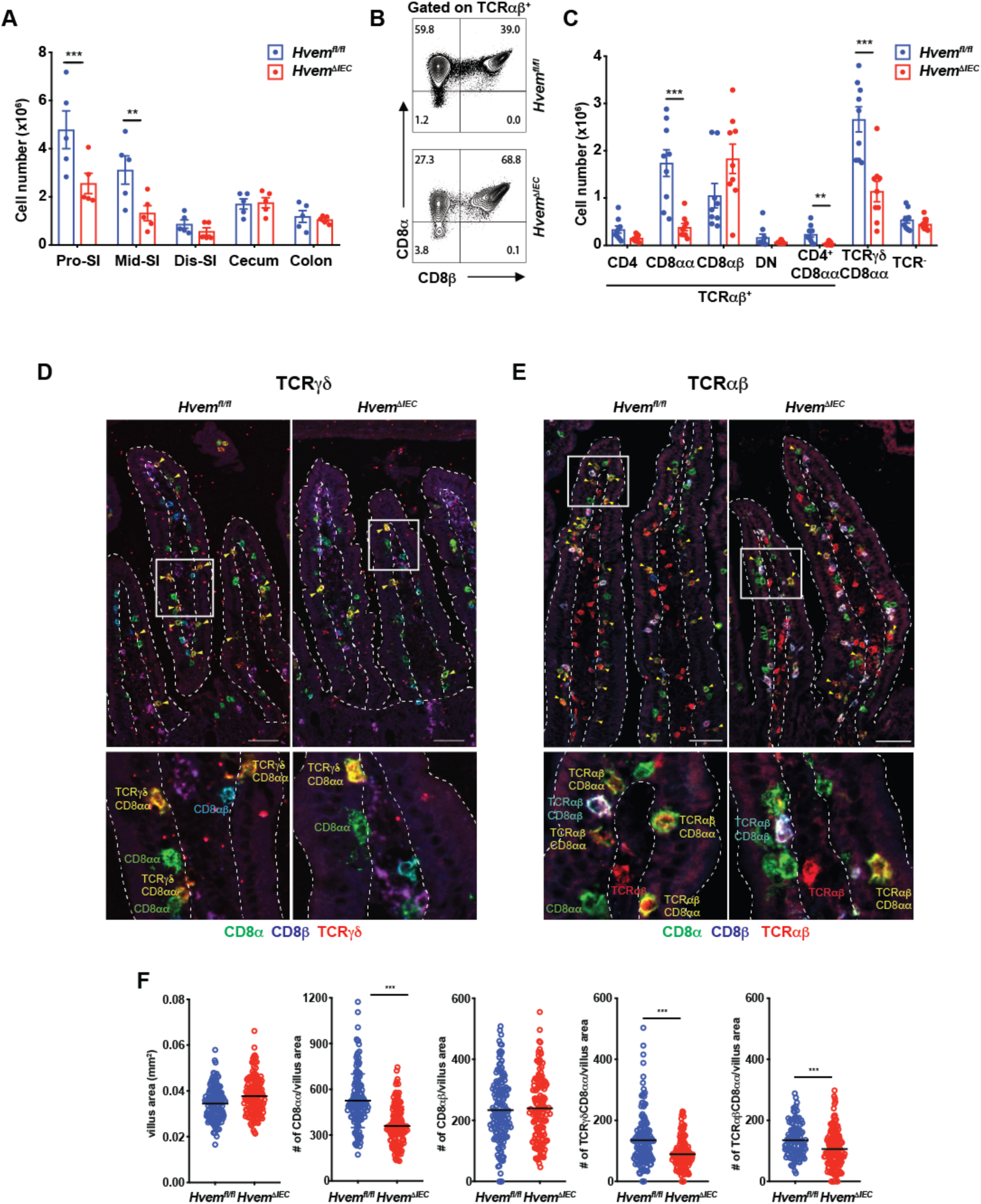
Epithelial HVEM is required for CD8αα^+^ IET. (A) Total IEL numbers in portions of the intestine from *Hvem*^*fl/fl*^ and *Hvem*^*ΔIEC*^ mice. (B) Representative plots of TCRαβ^+^CD8αα^+^ and TCRαβ^+^CD8αβ^+^ in TCRαβ IET from proximal SI in *Hvem*^*fl/fl*^ and *Hvem*^*ΔIEC*^ mice. (C) Absolute numbers of indicated IET subsets in total IEL from proximal SI in *Hvem*^*fl/fl*^ and *Hvem*^*ΔIEC*^ mice. (D-E) Representative immunofluorescence staining of TCRγδ IET (D) and TCRαβ IET (E) from proximal SI in *Hvem*^*fl/fl*^ and *Hvem*^*ΔIEC*^ mice. Composite images in which the three channels were merged. Composite images depict expression of CD8α (green), CD8β (blue), and TCRδ (red, D) or TCRβ (red, E). Yellow arrowheads highlight TCRγδ^+^CD8αα^+^ (D) and TCRαβ^+^CD8αβ^+^ IET (E). Single-channel images are in Figure S3. Dashed white lines indicate the boundaries of the epithelium, approximately one villus in each case, used for the quantitation. Scale bars, 50μm. Two independent experiments were carried out yielding similar results. (F) Quantification of TCRγδ^+^CD8αα^+^ and TCRαβ^+^CD8αβ^+^ per mm^2^ from proximal SI in *Hvem*^*fl/fl*^ and *Hvem*^*ΔIEC*^ mice. Statistical analysis was performed using unpaired t-test (A, C, F). **, p < 0.01; ***, p < 0.001. In A and C, bars show the mean and each symbol represents a measurement from a single mouse. In F, lines show the mean and each symbol represents a calculation to give the number per mm^2^. Data are representative results from one of at least two independent experiments with at least four mice in each experimental group (A) or pooled results from at least two independent experiments with at least four mice per group in each experiment (C, F). Groups of co-housed littermates were analyzed. See also Figures S2 and S3.

To confirm that the homeostasis of IET in the small intestine is affected by HVEM expression, we performed *in situ* immunofluorescence (IF) multi-color staining in frozen sections from the proximal small intestine of *Hvem*^*ΔIEC*^ and control mice. To identify TCRγδ^+^CD8αα^+^ IET, we used fluorochrome-coupled antibodies to CD8α, CD8β and TCRδ to detect CD8α^+^CD8β^-^TCRδ^+^ cells (Figure S3A). Similarly, to identify TCRαβ^+^CD8αα^+^ IET, we used fluorochrome-coupled antibodies to CD8α, CD8β and TCRβ to identify CD8α^+^CD8β^-^TCRβ^+^ cells (Figure S3B). Although this method did not distinguish TCRαβ^+^CD8αα^+^ IET from the subset of CD4^+^ CTL with a TCRαβ^+^CD4^+^CD8αα^+^ phenotype, the CD4^+^ CTL were much less frequent. After imaging by confocal microscopy, the IET subsets were quantified (Figures 2D-2F). The total villus area in the proximal SI was comparable between *Hvem*^*fl/fl*^ and *Hvem*^*ΔIEC*^ mice (Figure 2F). Similar to the flow cytometry results, *Hvem*^*ΔIEC*^ mice had a decrease of CD8αα^+^ IET including TCRαβ^+^CD8αα^+^ IET and TCRγδ^+^CD8αα^+^ IET (Figures 2D-2F). Although the number of CD8αβ^+^ IET was decreased by flow cytometry in *Hvem*^*-/-*^ mice, this was not observed in the proximal SI of *Hvem*^*ΔIEC*^ by flow cytometry or by confocal microscopy (Figures 2C-2F) and therefore the less pronounced decrease TCRαβ^+^CD8αβ^+^ IET could reflect HVEM action in another cell type.

### Epithelial HVEM is required for natural IET survival

Among the possible mechanism(s) for the decreased number of IET in *Hvem*^*ΔIEC*^ mice, we tested for components involved in homing to the intestine or the maintenance of the cells in the epithelial site, IET proliferation and IET survival. The homing and maintenance of intestinal IET in the epithelium depends on expression of β_7_ integrins, including α4β7 and the α_E_β_7_ integrin, which interacts with E-cadherin expressed by IEC (Cepek et al., 1994; Higgins et al., 1998; Schon et al., 1999). Flow cytometric analysis indicated that the expression of the integrin β7 chain and the integrin αE were not affected in IET from *Hvem*^*ΔIEC*^ mice (Figures S4A, S4B). The expression of mRNA for the *Cdh1* gene encoding E-cadherin in IEC was normal as well (Figure S4C).

To address if proliferation was involved, Ki-67 staining and EdU incorporation assays were performed. Although the trend suggested increased Ki-67, the IET from *Hvem*^*ΔIEC*^ mice were not significantly different from the controls (Figure S4D). Similarly, after six days of continuous EdU labeling, the percent of labeled IET was not different in *Hvem*^*ΔIEC*^ mice. In an experiment in which the EdU label was chased or removed for 14 days, however, terminating EdU administration led to a decreased percentage in EdU^+^ IET subsets compared to control mice (Figure 3A). These data suggested there was a more prominent survival defect in the TCRαβ^+^CD8αα^+^ and TCRγδ^+^CD8αα^+^ IET in *Hvem*^*ΔIEC*^ mice. Compared with control mice, cells from *Hvem*^*ΔIEC*^ mice exhibited higher Annexin V^+^ labeling in both the TCRαβ^+^CD8αα^+^ and TCRγδ^+^CD8αα^+^ IET subsets when analyzed ex vivo (Figures 3B, S4E). They also expressed greater amounts of the pro-apoptotic Bcl2 family proteins Bim and Bax (Figures S4F, S4G). To confirm that the absence of epithelial HVEM affected IET survival, we analyzed mice with a TCRαβ cell-specific overexpression of the pro-survival Bcl2 family gene *Bclxl* (*Bcl2l1*) driven by the *Lck* proximal promoter (*Lck*^*pr*^*-Bcl-xL*^*Tg*^) crossed to either *Hvem*^*fl/fl*^ or *Hvem*^*ΔIEC*^ mice. *Hvem*^*ΔIEC*^ mice expressing the *Bclxl* transgene had a normal number of TCRαβ^+^CD8αα^+^ IET. The increase in TCRγδ^+^CD8αα^+^ IET was not significant (Figure 3C), but the proximal Lck promoter does not work efficiently in TCRγδ^+^ cells (Fiala et al., 2019). Overall, the data are consistent with the hypothesis that epithelial HVEM affects the survival of natural IET populations, most prominently TCRαβ^+^CD8αα^+^ IET. To gain further insight into the effects of epithelial HVEM on TCRαβ^+^CD8αα^+^ IET, we performed RNA-seq analysis on TCRαβ^+^CD8αα^+^ IET sorted from SI of *Hvem*^*ΔIEC*^ and co-housed littermate *Hvem*^*fl/fl*^ mice. The top 50 differentially expressed genes contained 19 genes that have been reported to affect cell cycle, cell survival or cell growth (Figures 3D, 3E). Among those genes reduced in IET from *Hvem*^*ΔIEC*^ mice were autophagy and beclin 1 regulator 1 (*Ambr1*), which promotes cell survival under stress conditions by inducing autophagy (Fimia et al., 2013), cullin associated and neddylation dissociated gene 1 (*Cand1*), with reduced expression associated with apoptosis in cancer cells (Che et al., 2018) and also, there was decreased expression of some genes in the NF-κB pathway.

**Figure 3.**
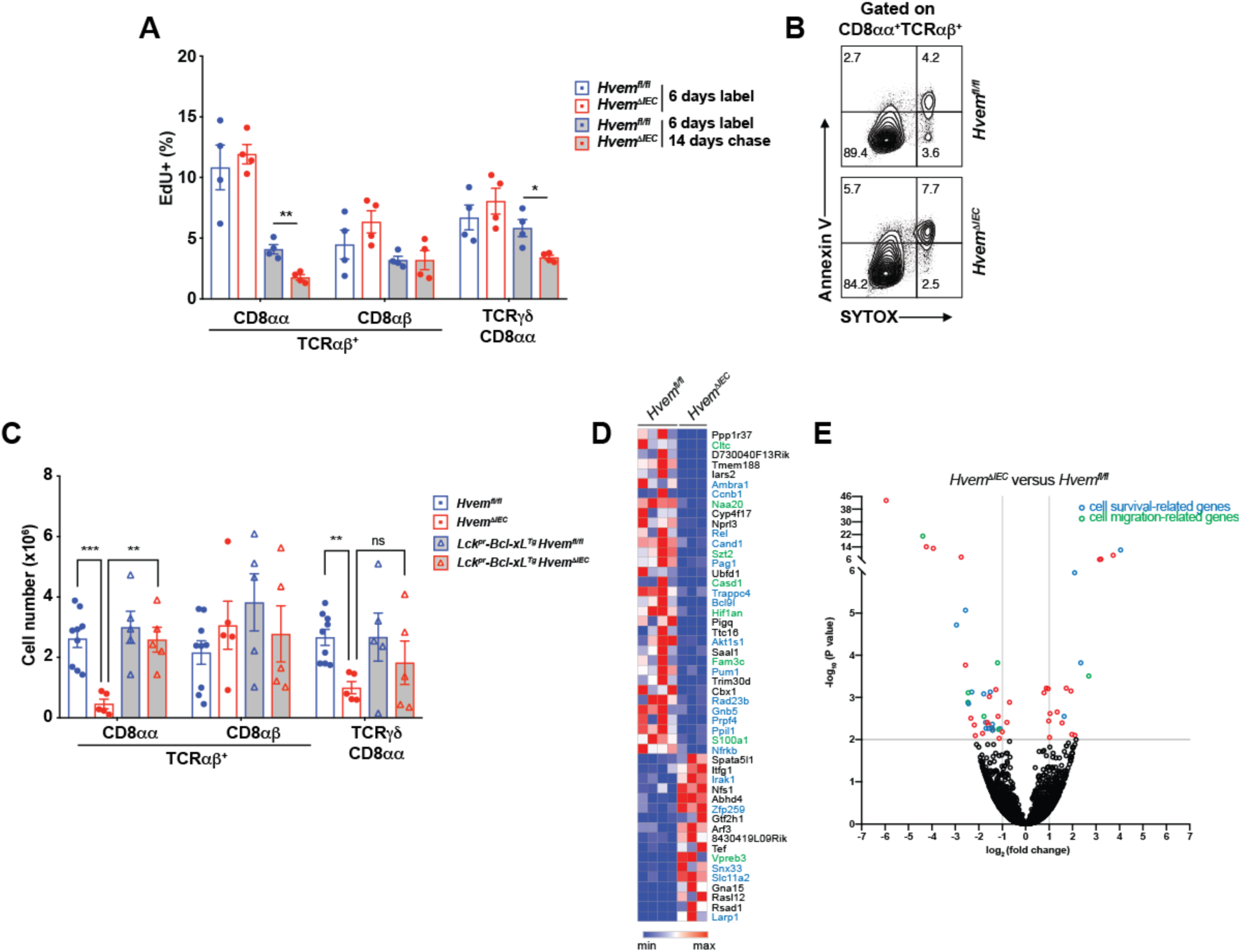
Epithelial HVEM affects CD8αα^+^ IET survival. (A) Frequencies of EdU ^+^ cells from proximal SI IET of *Hvem*^*fl/fl*^ and *Hvem*^*ΔIEC*^ mice. Mice were administered EdU once per day for 6 days. Cells were isolated from proximal SI and EdU^+^ cells were measured at either day 6 or day 20. (B) Representative flow cytometry images detecting apoptotic cells in TCRαβ^+^CD8αα^+^ IET from proximal SI in *Hvem*^*fl/fl*^ and *Hvem*^*ΔIEC*^ mice. (C) Absolute cell number of TCRαβ^+^CD8αα^+^, TCRαβ^+^CD8αβ^+^, and TCRγδ^+^CD8αα^+^ IET subsets from *Hvem*^fl/fl^, *Hvem*^*ΔIEC*^, *Lck*^*pr*^*-Bcl-xL*^*Tg*^*Hvem*^*fl/fl*^ and *Lck*^*pr*^*-Bcl-xL*^*Tg*^*Hvem*^*ΔIEC*^ mice. Statistical analysis was performed using an unpaired t-test. Statistical significance is indicated by *, p < 0.05; **, p < 0.01; ***, p < 0.001; ns, not significant. Bars show the mean and each symbol represents a measurement from a single mouse. Data are representative results of one of at least two independent experiments with at least four mice in each experimental group. (D) Sorted TCRαβ^+^CD8αα^+^ IET from SI of *Hvem*^*fl/fl*^ and *Hvem*^*ΔIEC*^ mice were analyzed by RNA-seq. The top 50 most differentially expressed genes with respect to *P*-value. (E) Volcano plots showing mean log_2_-transformed fold change (x axis) and significance (-log_10_ (adjusted P value)) of differentially expressed genes between the TCRαβ^+^CD8αα^+^ IET from the SI of *Hvem*^*fl/fl*^ and *Hvem*^*ΔIEC*^ mice. In D and E, genes in blue font or blue symbol are associated with cell survival or proliferation and those designated with green are associated with cell migration. Groups of co-housed littermates were analyzed. See also Figure S4.

### HVEM affects IET patrolling

The top 50 differentially expressed genes in IET from wild type and *Hvem*^*ΔIEC*^ mice also contained eight genes that have been reported to influence cell migration (Figures 3E, 3F). Included in these genes with decreased expression in *Hvem*^*ΔIEC*^ mice were clathrin heavy chain (*Cltc*), which among its different functions influences lymphocyte migration by inducing actin accumulation at the cellular leading edge (Ramirez-Santiago et al., 2016), N-α-acetyltransferase 20 (*Naa20*), which acetylates some terminal methionine amino acids and affects the actomyosin fibers needed for cell migration (Van Damme et al., 2012), and CAS1 domain containing protein 1 (*Casd1*), which catalyzes the 9-O-acetylation of sialic acids and which is required for cell migration at least in the nervous system (Santiago et al., 2004).

To further examine how epithelial HVEM deficiency affected IET, and particularly the migration of these cells, we used intravital imaging to track lymphocytes labeled with a CD8α-specific mAb in the epithelium and lamina propria of the SI in live mice (Edelblum et al., 2012; Wang et al., 2014). We found that the average coverage of the villus area by migrating CD8α labelled cells was greatly reduced in *Hvem*^*ΔIEC*^ mice (Figures 4A-4C). This decrease in coverage reflected not only the decreased cell number described above, but also average decreases in IET movement. The mean speed (Figures 4D) (Movies S1 and S2), track length (Figures 4E) and displacement (Figures 4F) of CD8α mAb-labeled IET were all decreased in *Hvem*^*ΔIEC*^ mice. The CD8α labelling detects both CD8αβ^+^ and CD4^+^CD8α^+^ IET as well as some ILC1, therefore it is possible that their movements were also affected. These data indicate that epithelial HVEM expression is required not only for the accumulation of normal population of subpopulations of IET, but also for the patrolling behavior of CD8α expressing cells.

**Figure 4.**
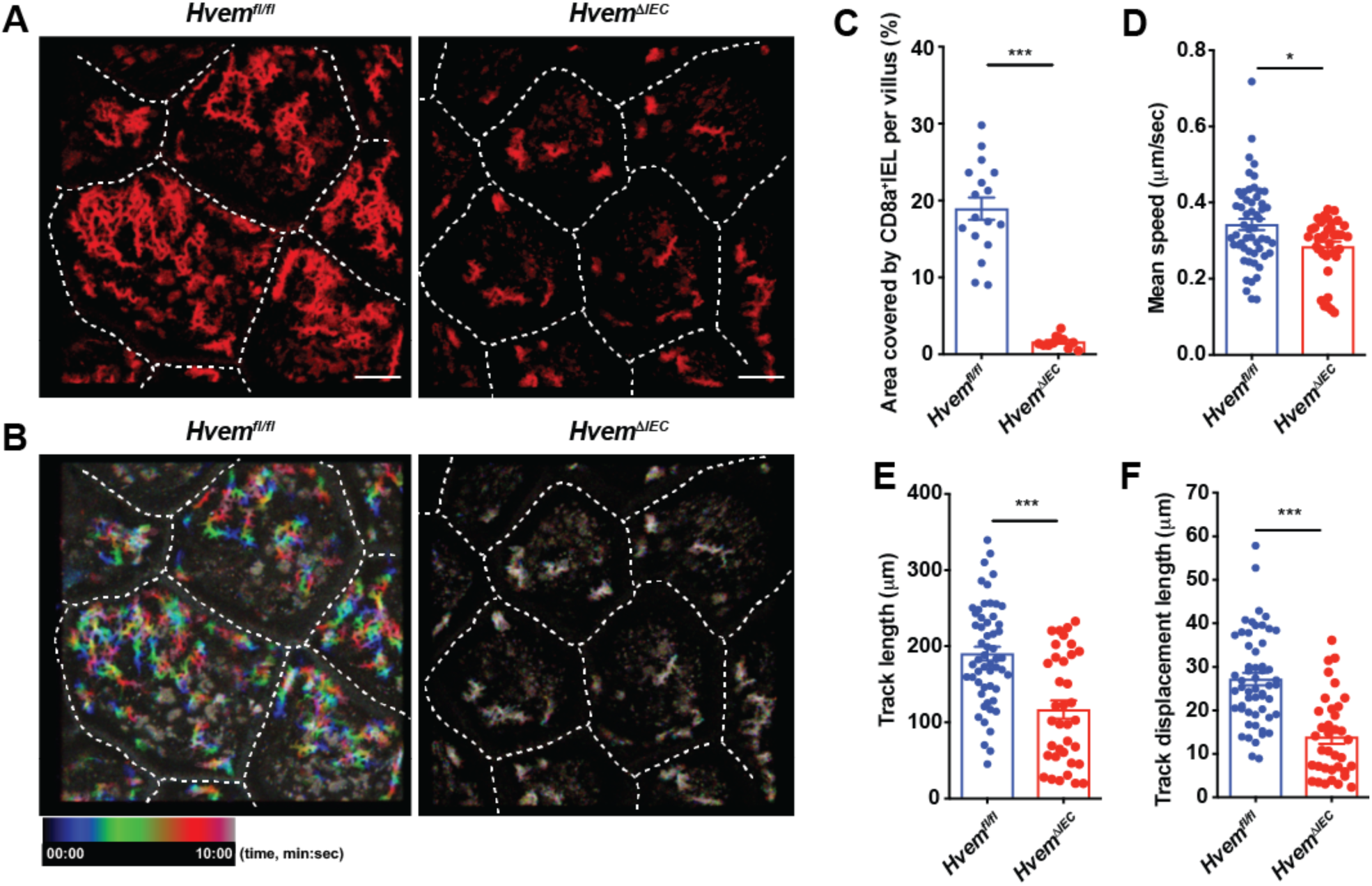
Epithelial HVEM influences patrolling by CD8α^+^ IET. (A) Time projection reveals the area covered by CD8α^+^ IET (red) during 10 minutes. Scale bars, 50μm. (B) Representative image of time-coded tracks made by CD8α^+^ IET cells in 10 minutes. Blue denotes starting position at the beginning of the recording, red denotes ending position. Rainbow color shows the tracks of moving CD8α^+^ IET. Quantification of the area covered by CD8α^+^ cells (C), mean speed (D), length of entire path or track length (E), and the distance between beginning and ending positions or displacement (F) in a 10 minute recording. Statistical analysis was performed using Mann-Whitney test (C-F). Statistical significance is indicated by *, p < 0.05; ***, p < 0.001. C-F, bars show the mean and each symbol represents a measurement from a single cell. Dashed white lines (A, B) indicate the boundaries of each quantified villus. Data represent combined results of two independent experiments with at least three mice in each experimental group. Groups of co-housed littermates were analyzed. See also supplemental movies S1and S2.

### HVEM signaling increases basement membrane synthesis

To interpret how epithelial HVEM affects CD8αα^+^ IET, especially IET survival, we performed RNA-seq analysis on IEC sorted from proximal SI of *Hvem*^*ΔIEC*^ and co-housed littermate *Hvem*^*fl/fl*^ mice. By gene set enrichment analysis, we observed a significant down-regulation of a number of genes associated with the extracellular matrix in *Hvem*^*ΔIEC*^ mice (Figure S5A). Down-regulated genes in IEC from *Hvem*^*ΔIEC*^ mice revealed significant enrichment in genes associated with the proteinaceous extracellular matrix according to gene ontology (GO) terms (Figure 5A). In agreement with this, analysis of differentially expressed genes also identified extracellular matrix, basement membrane and collagen, all categories decreased in the absence of epithelial HVEM expression (Figure S5B, S5C). These results suggest a possible role for epithelial HVEM in supporting IET survival and migration through stimulation of production of extracellular matrix proteins.

**Figure 5.**
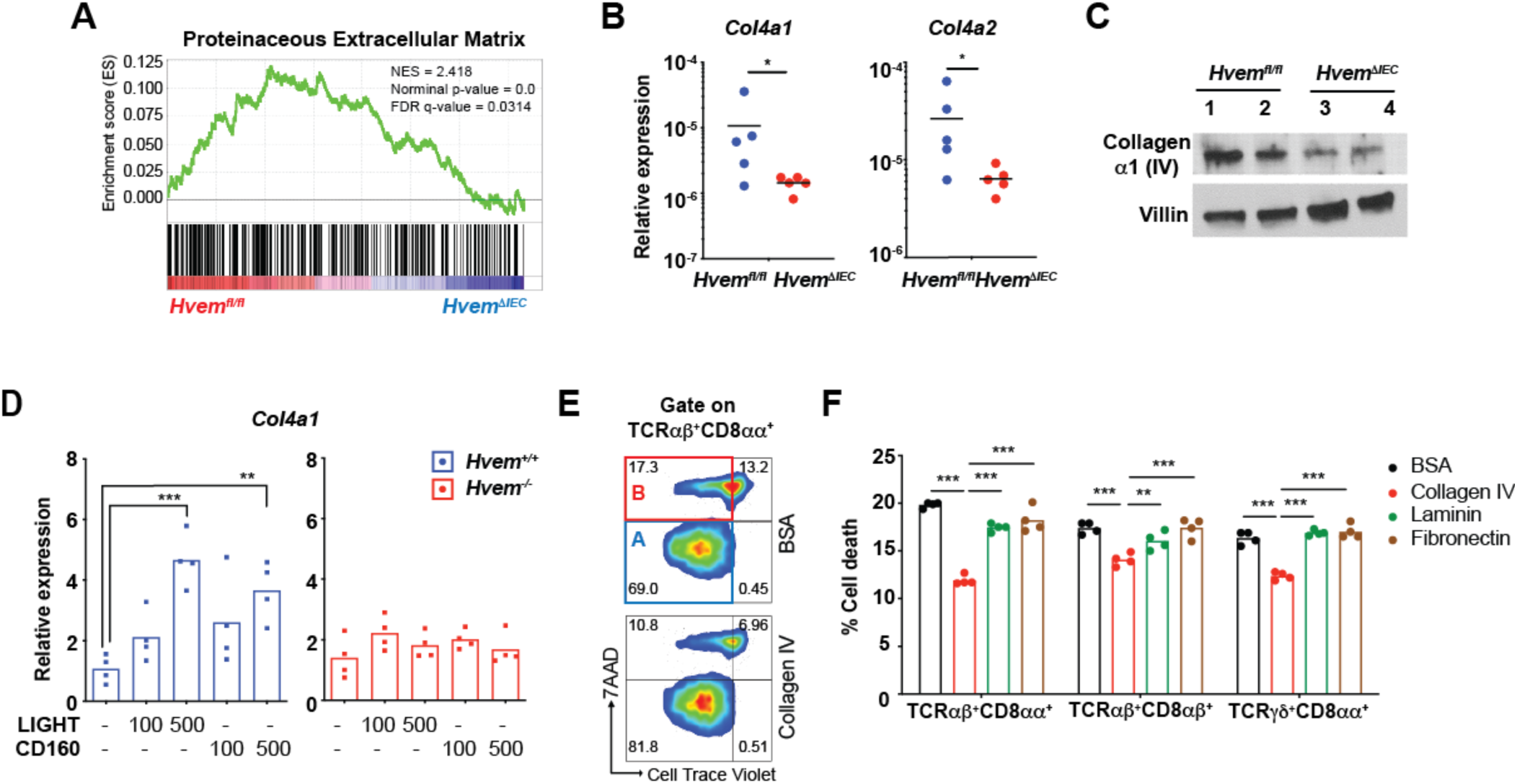
HVEM signaling contributes to induction of basement membrane synthesis. (A) Gene set enrichment analysis (GSEA) of transcripts from isolated IEC (CD31^-^CD45^-^EpCAM^+^) showing downregulation of GO Proteinaceous Extracellular Matrix genes in *Hvem*^*ΔIEC*^ mice. (B) Gene expression of *Col4a1* and *Col4a2* mRNA in IEC from *Hvem*^*fl/fl*^ and *Hvem*^*ΔIEC*^ mice by q-PCR. Data are normalized to the *Actb* housekeeping gene. (C) Expression of collagen α1 (IV) in sorted IEC from *Hvem*^*fl/fl*^ and *Hvem*^*ΔIEC*^ mice by Western blot. The expression level of villin is used as a control for the amount of protein lysates loaded. (D) Gene expression of *Col4a1* mRNA determined by q-PCR in cells from intestinal organoid cultures from proximal SI of *Hvem*^*fl/fl*^ and *Hvem*^*ΔIEC*^ mice. Isolated crypts from *Hvem*^*+/+*^ and *Hvem*^*-/-*^ mice were cultured with growth factors in the presence or absence of HVEM ligands, indicated concentrations in ng/ml, for 7 days. Data are normalized to *Rpl32* as the housekeeping gene. (E-F) Isolated SI CD8α^+^ IET (CD4^-^CD19^-^EpCAM^-^Ter^-^119^-^CD8α^+^) from wild type mice were labeled with CellTrace™ Violet (CTV) and cultured on plates coated with the indicated mouse basement proteins or BSA control. After 3 days, cells were stained with antibodies and 7AAD for monitoring cell death by flow cytometry. Representative plots (E) and calculated of % cell death (F) in TCRαβ^+^CD8αα^+^ IET. Statistical analysis was performed using an unpaired t-test. Statistical significance is indicated by *, p < 0.05; **, p < 0.01; ***, p < 0.001. Bars or lines show the mean and each symbol represents a measurement from a single mouse (B), single organoid culture (D) or an individual well (E, F). Data are representative results from one of at least two independent experiments. Groups of co-housed littermates were analyzed. See also Figure S5.

Collagen IV is a major component of basement membranes. Six type IV collagen alpha chains assemble into three types of heterotrimers, α1α1α2 (2:1 ratio), α3α4α5, and α5α5α6 (Mao et al., 2015). Collagen composed of the α1 (IV) and α2 (IV) chains is present in nearly all basement membranes (Timpl, 1989), while the other types have more restricted distributions. To further test if epithelial HVEM regulates the production of collagen IV in vivo, we carried out quantitative real-time PCR (qPCR) and showed reduced expression of *Col4a1* and *Col4a2*, mRNA, encoding the collagen α1 (IV) and α2 (IV) chains, respectively, by IEC from *Hvem*^*ΔIEC*^ mice (Figure 5B). *Hvem*^*ΔIEC*^ IEC also exhibited reduced collagen α1 (IV) protein expression by Western blot (Figure 5C). Therefore, these results suggest a role for epithelial HVEM in contributing to the production of collagen IV in the proximal SI.

To address if HVEM signaling induces *Col4a1* mRNA expression in IEC, intestinal organoid cultures from *Hvem*^*ΔIEC*^ and control mice were stimulated *in vitro* with HVEM ligands, LIGHT or CD160. *Col4a1* mRNA expression was increased by either soluble LIGHT or CD160 in the wild type organoid culture, but not in *Hvem*^*ΔIEC*^ organoids (Figure 5D). Together, these data indicate that epithelial HVEM signals to stimulate increased synthesis of mRNA for collagen IV.

To investigate the role of the collagen IV and other basement membrane proteins, such as laminin and fibronectin, in promoting IET survival, we carried out an in vitro culture system. Isolated CD8α^+^ IET were cultured on plates coated with basement membrane proteins and cell death was monitored by flow cytometry. CD8αα^+^ natural IET populations cultured on collagen-coated wells had decreased cell death compared to CD8αα^+^ IET cultured on either laminin, fibronectin or BSA (Figures 5E, 5F). In this in vitro system, death of TCRαβ^+^CD8αβ^+^ was also decreased.

### Collagen-binding integrins increase IET survival

Integrins containing the β1 chain are known to bind collagen IV, especially α1β1 integrins, with α2β1 integrins binding to a lesser extent (Roberts et al., 1999). TCRαβ^+^CD8αα^+^ and TCRγδ^+^CD8αα^+^ IET express relatively high amounts of integrin α1 and β1 with lower amounts expressed by total CD4^+^ IET, in agreement with a previous report (Figure 6A) (Marsal et al., 2005). To investigate the role of integrins in IET survival, we blocked α1β1 integrin binding to collagen IV with a β1-specific antibody. This led to increased TCRαβ^+^CD8αα^+^ cell death in vitro (Figure 6B). Cell death was also increased using the peptide disintegrin (inhibitor) Obtustatin (Figure 6B). These data suggest that collagen IV might influence CD8αα^+^ IET survival in part through binding to α1β1 integrin.

**Figure 6.**
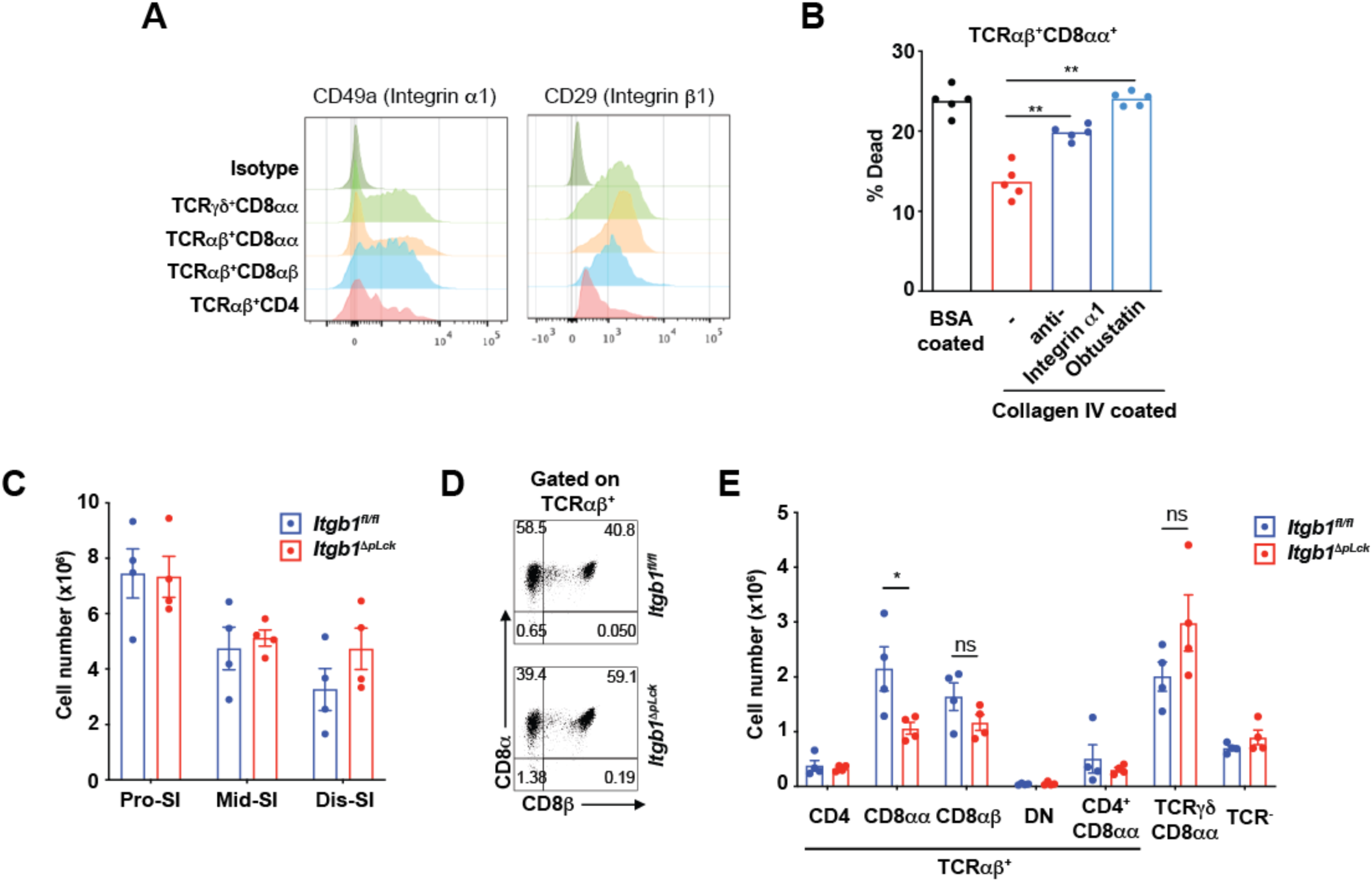
Collagen-binding integrins increase CD8αα^+^ IET survival. (A) Expression of integrin α1 and β1 subunits by IET populations from proximal SI of wild type mice by flow cytometry. (B) Isolated CD8α^+^ IET from SI of wild type mice labeled with CellTrace™ Violet (CTV) and cultured on plates coated with BSA control or collagen IV in the presence or absence of anti-integrin α_1_ mAb or inhibitor Obtusatin. After 3 days, cells were monitored for viability by flow cytometry. (C) Total IEL numbers in segments of SI from *Itgb1*^*fl/fl*^ and *Itgb1*^*ΔpLck*^ mice. (D) Representative plots of TCRαβ^+^CD8αα^+^ and TCRαβ^+^CD8αβ^+^ among TCRαβ IET from proximal SI in *Itgb1*^*fl/fl*^ and *Itgb1*^*ΔpLck*^ mice. (E) Absolute numbers of indicated subsets in total IEL from proximal SI in *Itgb1*^*fl/fl*^ and *Itgb1*^*ΔpLck*^ mice. Statistical analysis was performed using unpaired t-test. Statistical significance is indicated by *, p < 0.05; **, p < 0.01; ns, not significant. Bars show the mean and each symbol represents a measurement from an individual well (B) or a single mouse (C, E). Data are representative results from one of at least two independent experiments. Groups of co-housed littermates were analyzed.

To determine if there was a role for T cell-expressed integrins in CD8αα^+^ IET survival in vivo, we generated *Itgb1*^*ΔpLck*^ mice by crossing *Lck*^*pr*^-Cre mice with *Integrin b1*^*fl/fl*^ (*Itgb* ^*fl/fl*^) mice. Flow cytometric analyses indicated that *Itgb1*^*ΔpLck*^ mice had no difference in total CD45^+^ cell numbers throughout SI compared to *Itgb1*^*fl/fl*^ controls (Figure 6C). However, the TCRαβ^+^CD8αα^+^ IET subset in the proximal SI of *Itgb1*^*ΔpLck*^ mice was decreased (Figures 6D, 6E), consistent with the greater activity of the *Lck*^*pr*^*-*Cre in TCRαβ cells (Fiala et al., 2019). Therefore, this selective effect is consistent with the hypothesis that basement membrane collagen IV promotes IET survival, likely in part through α_1_β_1_ integrin binding to collagen IV.

## Discussion

IET are tissue-resident T lymphocytes in close contact with the epithelium of the intestine. Maintenance and function of these tissue resident lymphocytes likely requires complex sets of signals from the microbiome, tissue cells including the epithelium, and extracellular matrix. The mechanisms dependent on IEC that regulate IET homeostasis in the small intestine are not well understood. Here, we provide evidence that HVEM, a TNFR superfamily member that is constitutively expressed by IEC, is involved in the survival of IET subsets. Furthermore, epithelial HVEM was important for normal migration of IET and patrolling of the villi in the small intestine. HVEM does this in part by signaling to the IEC, leading to increased expression of components of the basement membrane, such as collagen IV. Collagen IV interacts with IET integrins and it promoted the survival of IET populations in vitro via binding to β1 integrins. Moreover, TCRαβ^+^CD8αα^+^ IET were reduced in *Itgb1*^*ΔpLck*^ mice. Therefore, our data have defined a novel circuit for tissue resident lymphocyte homeostasis and function in the intestine whereby signals to epithelial cells communicate through extracellular matrix synthesis to resident T lymphocytes.

The effects of epithelial HVEM deficiency on survival were not uniform across different IET populations. The presence of TCRαβ^+^CD8αα^+^ IET in the small intestine were most consistently and severely reduced. Another large population of natural IET, TCRγδ^+^ IET, were also decreased in most assays. Induced, TCRαβ^+^ CD8αβ^+^ IET were not decreased in *Hvem*^*ΔIEC*^ mice, although these IET express α1β1 integrins and had enhanced survival in vitro in the presence of collagen IV. This population of induced IET was not dependent on β1 integrins in vivo and likely makes use of additional survival mechanisms or alternatively, it may be replenished more continuously by CD8^+^ T cells primed in mesenteric lymph nodes. Although small intestine IET depend on β7 integrins and CCR9 for homing and cell adhesion (Cepek et al., 1994; Staton et al., 2006), There are precedents for signals that selectively influence IET subsets. For example, *AhR*-deficient mice had reduced survival only of natural IET (Li et al., 2011). IL-15-IL-15R interactions also most severely affected the natural IET subsets (Hu et al., 2018; Lodolce et al., 1998; Ma et al., 2009). TCRγδ^+^ IET unlike the others subsets, were not highly dependent on the microbiota (Bandeira et al., 1990; Umesaki et al., 1993), but they were influenced by expression of GPR18 and GPR55 (Wang et al., 2014) (Sumida et al., 2017). Furthermore, epithelial expression of the thymus leukemia (TL) antigen, a nonclassical class I molecule, shapes the population of induced TCRαβ^+^ CD8αβ^+^ IET via their ability to induce co-expression of CD8αα homodimers (Huang et al., 2011). Therefore, some degree of variability of the effect on IET populations in *Hvem*^*ΔIEC*^ mice was not surprising.

IET are migratory cells located above the basement membrane, which they patrol extensively (Edelblum et al., 2012). Movement and survival of IET may be linked under some circumstances, as both were negatively regulated in TCRγδ^+^ IET by GPR55 (Sumida et al., 2017). RNA-Seq analyses of confirmed that the expression of genes related to survival and genes related to migration were reduced in TCRαβ^+^CD8αα^+^ IET in the absence of epithelial HVEM. Consistent with this, in *Hvem*^*ΔIEC*^ mice IET speed, track length and displacement were all decreased, leading to a greatly decreased area of the villi covered. These effects were evident when all CD8α^+^ IET were labeled, suggesting there is perhaps a broader effect on IET subsets in addition to TCRαβ^+^CD8αα^+^ IET. IET movement and surveillance may be important for protecting the epithelium from injury or infection. Previously it has been shown that TCRγδ^+^ IET movement in the small intestine is dynamically regulated by MyD88 dependent signaling by IEC (Hoytema van Konijnenburg et al., 2017). Furthermore, exposure to invasive *Salmonella* bacteria enhanced the movement of TCRγδ^+^ IET between the IEC in a “flossing” movement.

Based on RNA-Seq of IEC and other analyses, our data suggested decreased extracellular matrix protein synthesis by epithelial cells from *Hvem*^*ΔIEC*^ mice. Signaling by HVEM ligands in vitro increased *Col4a1* and *Col4a2* mRNA synthesis in intestinal organoid cultures, providing direct evidence connecting HVEM signaling in IEC to increased type IV collagen synthesis. Furthermore, microwells coated with type IV collagen promoted IET survival in vitro, providing a link between HVEM signals and IET survival. This effect of collagen IV is consistent with previous reports showing that it promotes survival and proliferation of different types of cancer cells (Burnier et al., 2011; Ohlund et al., 2013). Collagen IV also was shown to promote migration of tumor cells and endothelial cells (Chelberg et al., 1989; Herbst et al., 1988). Therefore, epithelial HVEM deficiency likely affected the patrolling of IET in part through altered basement membrane synthesis. The effects of increasing type IV collagen are described here, but our data do not exclude HVEM-dependent effects IET mediated by other extracellular matrix proteins.

Various integrins containing a β_1_ subunit bind to extracellular membrane components, including different types of collagen and fibronectin (Barczyk et al., 2010). For example, integrin α1β1 preferentially binds collagen type IV, while α_2_β_1_ preferentially binds to collagen type I (Barczyk et al., 2010; Eble et al., 1993; Roberts et al., 1999). It has been shown that human IEL bind preferentially to collagen VI as opposed to laminin and fibronectin through integrin α1β1 (Roberts et al., 1999). Mouse IEL also bind collagen (Marsal et al., 2005), and we confirmed that IET express collagen-binding integrins. The positive survival effect of TCRαβ^+^CD8αα^+^ IET culture on collagen IV coated microwells could be reversed by blocking integrin binding. Moreover, mice with a deficiency for *Itgb1* in TCR αβ^+^ cells had reduced TCRαβ^+^CD8αα^+^ IET. Previous in vivo analyses of the role of α_1_β_1_ integrins provided conflicting results on their importance for IET accumulation (Marsal et al., 2005; Meharra et al., 2000). The designs of the previous experiments were very different, but regardless, in neither study were the TCRαβ^+^CD8αα^+^ IET separately analyzed. We note that the overall effect of *Itgb1* deficiency in TCR αβ^+^ cells was milder than in *Hvem*^*ΔIEC*^ mice, consistent with the notion that there likely are other effects of epithelial HVEM deficiency on IET.

HVEM is a multi-functional protein that previously was shown to influence inflammation and the response to acute infections in the intestinal mucosae (Breloer et al., 2015; Seo et al., 2018; Shui et al., 2012; Steinberg et al., 2013). Here our data indicate that HVEM expression by IEC also has a crucial role in regulating mucosal immunity at steady state, providing a striking example of an intestinal epithelial cell surface receptor that affects both natural IET survival and IET migration. The putative functions of the natural IET populations include protection of barrier integrity and wound healing and responses to infectious agents (Hoytema van Konijnenburg et al., 2017; Nielsen et al., 2017; Saurer et al., 2004). Future experiments will help to unravel the extent to which the diverse effects of HVEM deficiency on colitis and mucosal infections are related to its effects on resident IET.

## Methods

### Mice

*Hvem*^*-/-*^ and control *Hvem*^*fl/fl*^ mice were bred and described previously (Seo et al., 2018). *Cd160*^*-/-*^ mice were provided by Dr. Yang-Xin Fu (UT Southwestern, TX) (Tu et al., 2015). *Cd4-*Cre (STOCK Tg(Cd4-cre)1Cwi/BfluJ; Stock No: 017336), *Villin-*Cre (B6.Cg-Tg(Vil1-cre)997Gum/J, Stock No: 004586), *Lck*^*pr*^-Cre (B6.Cg-Tg(Lck-cre)548Jxm/J, Stock No:003802), *Itgb1*^*fl/fl*^ (B6;129-*Itgb1*^*tm1Efu*^/J, Stock No: 004605) and *Lck*^*pr*^*-Bcl-xL*^*Tg*^ mice (B6.Cg-Tg(LCKprBCL2L1)12Sjk/J; Stock No: 013738) were all purchased from The Jackson Laboratory. *Lck*^*pr*^*-Bcl-xL*^*Tg*^ mice express the pro-survival Bcl2 family gene *Bclxl* (*Bcl2l1*) under the control of the mouse *Lck* proximal promoter (*Lck*^*pr*^ Cre). *Hvem*^*fl/fl*^ mice were bred to *Villin-cre* mice to generate conditional knockout mice, *Hvem*^*ΔIEC*^. *Integrin b1*^*fl/fl*^ mice were bred to *Lck*^*pr*^ Cre mice to generate conditional knockout mice, *Itgb1*^*ΔpLck*^ mice. All Cre mouse strains were maintained on the C57BL/6 background or were backcrossed to C57BL/6 for at least for 6 generations. 15 to 17-week-old male mice on B6 genetic background were used in this study. Whenever possible, groups of control and gene knockout mice were housed in the same cage to minimize the effect of housing conditions on experimental variation. For tissue or cell analyses, tissues were collected and used for immunofluorescence analysis and intraepithelial lymphocyte preparation. Mice were bred and housed under specific pathogen-free conditions in the vivarium of the La Jolla Institute for Immunology (LJI). All procedures were approved by the LJI Animal Care and Use Committee.

### Germ-free mouse husbandry

Germ-free mice were bred and housed in the animal facility of POSTECH Biotech Center. Germ-free C57BL/6 (B6) mice were kindly provided originally by Drs. Andrew Macpherson (Bern Univ., Switzerland) and David Artis (Univ. Pennsylvannia, USA) and maintained in sterile flexible film isolators (Class Biological Clean Ltd., USA). The sterility of germ-free mice was checked regularly by the absence of bacterial colonies in the culture experiment using their fecal pellets. Germ free C57BL/6 mice maintained in the animal facility of POSTECH Biotech Center in accordance with institutional ethical guideline and the protocols approved by the Institutional Animal Care and Use Committees (IACUC) of the POSTECH.

### Isolation of IEL

Intestines were collected from mice, and were divided into five parts, including proximal SI, middle SI, and distal SI, approximately one-third each, as well as cecum and colon. Peyer’s patches were carefully removed, and tissues were cut open longitudinally, briefly washed, and cut into 1.5 cm pieces. The tissue pieces were incubated in 20 mL of RPMI (5% FBS, 25 mM HEPES and 1 mM DTT) in a shaker at 200 rpm, 37°C, for 20 min, followed by incubated in 20 mL HBSS (25 mM HEPES and 20 mM EDTA) in a shaker at 200 rpm, 37°C, for 30 min. After each incubation, the cell suspension was filtered through a metal mesh and the supernatant was saved for IEL preparation. The flow-through cell suspension was spun down. The cell pellets were then re-suspended in 40% Percoll solution and overlaid above 80% Percoll solution carefully, followed by centrifugation at 2000 rpm, 25°C for 20 min without the brake. IEL were collected from the interface, washed once and re-suspended in the complete RPMI-1640 medium. These purified cells constituted the epithelial cell fraction enriched for IEL. The cells were used immediately for cell counting and staining.

### Flow cytometry

Flow cytometry analysis was performed on an LSRII instrument (BD Biosciences). The data were analyzed by using FlowJo software (Tree Star). Absolute cell counts were obtained by using CountBright™ Absolute Counting Beads (Life Technologies). The following mAbs were used: TCRδ (eBioscience, eBioGL3), TCRβ (BioLegend, H57-597), CD4 (eBioscience,RM4-5), CD8α (BD Biosciences, 53-6.7), CD8β (eBioscience, eBioH35-17.2), CD45 (BioLgegend, 30-F11), CD45.1 (BioLegend, A20), CD45.2 (Thermo Fisher Scientific, 104), integrin β7 (BD Biosciences, M293), integrin αE (Thermo Fisher Scientific, 2E7), Bax (Thermo Fisher Scientific, 6A7), Bim (Bio-Rad AbD Serotec, AHP933), CD49a (BD Biosciences, Ha31/8), CD29 (BioLegend, HMβ1-1), Ki-67 (Thermo Fisher Scientific, SP6), EdU (Thermo Fisher Scientific), Annexin V (BD Biosciences), SYTOX™ AADvanced™ Dead Cell Stain Kit (Thermo Fisher Scientific).

### Cryosection immunofluorescence

SI tissue was collected, opened longitudinally and the luminal contents washed in 20 mL RPMI in a 50 mL tube by inverting 20 times. The tissues were placed on foil with O.C.T. for the Swiss-roll technique. Once the entire intestine length was rolled up, the intestine Swiss roll was transferred into a tissue mold and was frozen on a thermo-conductive platform (ThermalTray, Biocision) pre-cooled with dry ice, and then placed into −80°C freezer. Frozen sections were cut to a 10 μm thickness. After drying at RT for 1 hour, frozen sections were fixed in pre-cooled ethanol at −20 °C for 10 min followed by pre-cooled acetone at −20 °C for 10 min. The fixed sections were used for immunofluorescence (IF) staining or kept in −80°C freezer for later use. Primary antibodies were diluted in PBST (PBS, 0.5% BSA, 0.1% Tween-20). Antibodies used were: CD45 (BioLegend, clone 30-F11), EpCAM (BioLegend, clone G8.8), TCRδ (BioLegend, clone GL3), TCRβ (BioLegend, clone H57-597), CD8α (BD, clone 53-6.7), CD8β (BD, clone H35-17.2). In Fig1B-1C, immunofluorescence imaging was performed on FV10i confocal microscope (Olympus). Areas from Pro-SI, Mid-SI or Dis-DI were outlined and CD45^+^ IEL per villus were counted manually. In Fig2D-2F, immunofluorescence imaging was performed on Zeiss LSM 780 or LSM 880 confocal microscopes (Zeiss). Data analysis was performed using ZEN software (Zeiss); areas of each villus from Pro-SI was outlined and measured and IET subsets of each villus were counted manually. The number of IET subsets in each villus was calculated in units of per mm^2^. ImageJ software (1.53c, National Institutes of Health)(Schindelin et al., 2012) were used for data for representative images.

### Intravital microscopy

Mice were injected with 15-20 µg of anti-CD8α-AF488 (eBioscience, clone 53-6.7) or anti-CD8α-AF647 (BD Biosciences, clone 53-6.7) with/without EpCAM-AF647 (BioLegend, clone G8.8) by retro-orbital injection 4 h before imaging. After anesthetization, mice were positioned on a WPI ATC2000 heating pad ventral side up and kept at 37°C. The distal duodenum was exposed and opened along the antimesenteric border. The mucosal surface was hydrated with PBS and placed against a coverslip fitted onto a suction ring (Looney et al., 2011). All intravital imaging was done using 25x 0.95NA water immersion objective on the Leica SP5 upright confocal microscope with a resonant scanner and acquired using Leica Application Suite. Images were acquired by taking Z-stacks encompassing the epithelium and the upper layers of the lamina propria at 2.8 um step size every 10 seconds. Each XY plane spans 365 × 365 µm^2^. Imaris software (version 9.2, Bitplane) or ImageJ software (1.53c, National Institutes of Health) were used for data processing and representative images and videos. Statistics (mean speed, track length, track displacement length) for each CD8α^+^ lymphocyte cell were calculated with Imaris and plotted using Prism (version 8, GraphPad).

### Small intestinal organoid culture

SI organoids were derived from *Hvem*^/-^ or littermate *Hvem*^+/+^ mice. The proximal SI was cut into 5-mm segments and incubated in 4°C 2 mM EDTA in PBS for 5 min and washed by pipetting. The segments were incubated in 2 mM EDTA in PBS for 30 min at 4°C, and crypts were isolated by pipetting with cold HBSS. Dissociated crypts were passed through a 70-μm cell strainer and pelleted by centrifuge at 600 rpm for 3 minutes at 4°C. The crypts were resuspended in Advanced DMEM/F12 medium (Thermo Fisher Scientific); the number of crypts was counted, and they were resuspended in Matrigel Growth Factor Reduced (GFR) Basement Membrane Matrix (Corning). The crypts were plated in a 24-well plate with organoid growth medium supplemented with 100 μg/ml penicillin, 100 U/ml streptomycin, 2 mM Glutamax, 1×N-2 supplement, 1×B27 supplement, 10 mM HEPES (Thermo Fisher Scientific), 1 mM N-acetylcysteine (Sigma-Aldrich), 100 ng/ml recombinant mouse Noggin (Peprotech), 50 ng/ml recombinant mouse EGF (BioLegend), 500 ng/ml recombinant human R-spondin 1 (Peprotech). Media were changed every 2 days.

### Quantitative Real-time PCR

Total RNA extraction from IECs was performed with RNeasy Kit (Qiagen), according to the manufacturer’s instructions. cDNA synthesis was performed by using iScript Advanced cDNA Synthesis Kit (Bio-Rad). Quantitative real-time PCR reactions were performed with SYBR Green I Master Kit and LightCycler 480 system (Roche). mRNA levels of *Col4a1* were normalized to the housekeeping gene *Actb or Rpl32*. The primers were synthesized by the Integrated DNA Technologies. Primers used were:

*Col4a1-forward 5’-TCCGGGAGAGATTGGTTTCC-3’*

*Col4a1-reverse 5’-CTGGCCTATAAGCCCTGGT-3’*

*Actb-reverse 5’-GATCTGGCACCACACCTTCT-3’*

*Actb - reverse 5’-GGGGTGTTGAAGGTCTCAAA-3’*

*Rpl32-reverse 5’-TTCCTGGTCCACAATGTCAA-3’*

*Rpl32 - reverse 5’-GGCTTTTCGGTTCTTAGAGGA-3’*

### RNA-seq library preparation

RNA-seq libraries were prepared from sorted proximal SI IEC or from sorted SI TCRαβ^+^CD8αα^+^ IET by the Sequencing Core at La Jolla Institute for Immunology using the SMARTer Stranded Total RNA-Seq Kit v2 - Pico Input Mammalian (TaKaRa)(Picelli et al., 2014). Libraries were sequenced on an Illumina HiSeq 2500, generating 50 bp single-end reads.

### RNA-seq analysis

The single-end reads that passed Illumina filters were filtered for reads aligning to tRNA, rRNA, adapter sequences, and spike-in controls. The reads were then aligned to mm10 reference genome using TopHat (v 1.4.1) (Trapnell et al., 2009). DUST scores were calculated with PRINSEQ Lite (v 0.20.3) (Schmieder and Edwards, 2011) and low-complexity reads (DUST > 4) were removed from the BAM files. The alignment results were parsed via the SAMtools (Li et al., 2009) to generate SAM files. Read counts to each genomic feature were obtained with the htseq-count program (v 0.7.1) (Anders et al., 2015) using the “union” option. After removing absent features (zero counts in all samples), the raw counts were then imported to R/Bioconductor package DESeq2 (v 1.6.3) (Love et al., 2014) to identify differentially expressed genes among samples. P-values for differential expression were calculated using the Wald test for differences between the base means of two conditions. These P-values were then adjusted for multiple test correction using the Benjamini-Hochberg algorithm (Benjamini and Hochberg, 1995). Principal Component Analysis (PCA) was performed using the ‘prcomp’ function in R. The sequences used in this article have been submitted to the Gene Expression Omnibus under accession number XXXX (http://www.ncbi.nlm.nih.gov/geo/).

Gene set enrichment analysis was done using the “GseaPreranked” method with “classic” scoring scheme. All the GO gene sets were downloaded from MSigDB in GMT format. Rank files for each DE comparison of interest were generated by assigning a rank of -log10(pValue) to genes with log2FoldChange greater than zero and a rank of log10(pValue) to genes with log2FoldChange less than zero. GO analysis of DEGs was performed with ToppGene (http://toppgene.cchmc.org)(Chen et al., 2009). FDR (Benjamini-Hochberg) < 0.05 was considered statistically significant. Heatmap was performed with Morpheus (https://software.broadinstitute.org/morpheus/). For classical pathway, Ingenuity Pathway Analysis (Qiagen) was used.

### Western-blot analysis

Harvested IECs were lysed in triton lysis buffer (137 mM NaCl, 20 mM Tris base at pH 7.4, 10% glycerol and 1% Triton X-100) supplemented with a protease and phosphatase inhibitor mixture (Roche) for 20min on ice, and centrifuged at 15,000 rpm for 10 minutes at 4°C. 10 μg of denatured proteins were loaded onto a Mini-PROTEAN Precast Gel (Bio-Rad) and transferred onto a PVDF membrane (Thermo Fisher Scientific).The membranes were blocked by 5% skim milk in TBST (Tris-buffered saline /0.1% Tween 20, Bio-Rad) and then incubated with antibodies to collagen a1 (IV) (Abcam). Immunoreactive bands were detected by chemiluminescence (ECL solution, Santa Cruz).

### CD8α^+^ IEL culture

For the isolation of IEL, CD8α^+^ cells were enriched from the SI IEL preparation by negative and positive selection with the iMag cell separation system (BD Biosciences) according to the manufacturer’s instructions. Briefly, single-cell IEL suspension was incubated with staining buffer (PBS containing 2% FBS with 2 mM EDTA) containing a mixture of biotin-conjugated mAbs against CD4 (RM4-5; BioLegend), CD19 (6D5; BioLegend), CD326 (G8.8; BioLegend), and TER-119 (TER-119; BioLegend) in a 5-ml round tube (BD Biosciences) for 30 minutes at 4°C. Cells were washed with staining buffer and incubated with Streptavidin Particle Plus-DM (BD Biosciences) for 30 minutes at 4°C. The tube was placed on the Cell Separation Magnet at room temperature for 8 minutes. The supernatant was carefully aspirated off as a negative fraction. The negative fraction was incubated with biotin-conjugated anti-CD8α (53-6.7; BioLegend) in staining buffer for 30 minutes at 4°C, washed with staining buffer, and incubated with Streptavidin Particle Plus-DM in a 5-ml round tube for 30 minutes at 4°C. The tube was placed on the Cell Separation Magnet at room temperature for 8 minutes. The supernatant was carefully removed and CD8α^+^ positive fraction was washed and resuspended with complete medium. CD8α^+^IEL were expanded in complete medium containing 50 ng/ml recombinant mouse IL-15/IL-15R complex (Thermo Fisher Scientific) on a tissue culture plate at the concentration of 5×10^5^ cells/ml for 3 days. CD8α^+^IEL were then cultured on a 96-well protein high binding plate (Corning) coated with 100 μg/ml bovine serum albumin (BSA, Thermo Fisher Scientific), type IV collagen (Corning), Laminin (Corning) or Fibronectin (Abcam). For monitoring cell death, isolated SI CD8α^+^ IET (EpCAM^-^CD4^-^CD19^-^TER119^-^ CD8α^+^) from wild type mice were labeled with CellTrace™ Violet (CTV, Thermo Fisher Scientific), and then labeled cells were incubated in the presence of recombinant IL-15/15Rα (50ng/mL) in the presence of absence of anti-integrin α1 (10μg/mL, BD Biosciences) or Integrin α1β1 inhibitor Obtusatin (10nM, R&D Systems) in the coated culture plate. After 3 days of culture, cells were stained with antibodies and 7-aminoactinomycin D (7AAD, BD Biosciences) for viability assessment of the different CD8α^+^ IET subsets when analyzed by flow cytometry. % cell death in proliferating cells is calculated as: % cell death in proliferating cells = B (CTV^-^7AAD^+^) / A (CTV^-^7AAD^-^) + B (CTV^-^7AAD^+^) x 100.

### Statistics

Details concerning the statistical analysis methods are provided in each figure legend. Briefly, all data were analyzed using GraphPad Prism 8 software and were shown as mean and the standard error of the mean (SEM). Statistical significance was determined by unpaired t-test for cell numbers, % of cell proliferation, cell survival and cell death and Mann-Whitney test for intravital microscopy. Statistical significance is indicated by *, p < 0.05; **, p < 0.01; ***, p < 0.001; ns, not significant.

## Supporting information

graphic abstract

highlights

## ACKNOWLEDGMENTS

This work was supported by grants from the National Institutes of Health (P01 DK46763, R01 AI61516 and MIST U01 AI125955 to M.K.; MIST U01 AI125957 to H.C; S10RR027366 to LJI Flow Cytometry Core Facility; S10OD021831 to LJI Microscopy Facility Core), the Crohn’s and Colitis Foundation of America (CCFA-254582 to Shui J-W), the Uehrara Foundation (to D.T.). We thank the staff of the Microscopy & Histology Core, Flow Cytometry Core, Sequencing Core, Bioinformatics Core and the Department of Laboratory Animal Care (DLAC) at La Jolla Institute for Immunology for excellent technical assistance, Christina Kim for RNA-seq library preparation, Sara MacArdel for help with the generation and analysis of microscopy data, and Ashu Sethi for analyzing RNA sequencing data.

## AUTHOR CONTRIBUTIONS

D.T., Q.W., G-Y.S. designed and performed the experiments and G-Y.S., Q.W. wrote the manuscript. D.T, Q.W., G-Y.S. Z.M., and P.M. performed the experiments and analyzed microscopy data. H.C. provided guidance and helped with writing the paper. All authors provided critical comments. M.K. supervised the study, designed experiments, and wrote the paper.

## Supplementary Figures

**Figure S1, Related to Figure 1.**
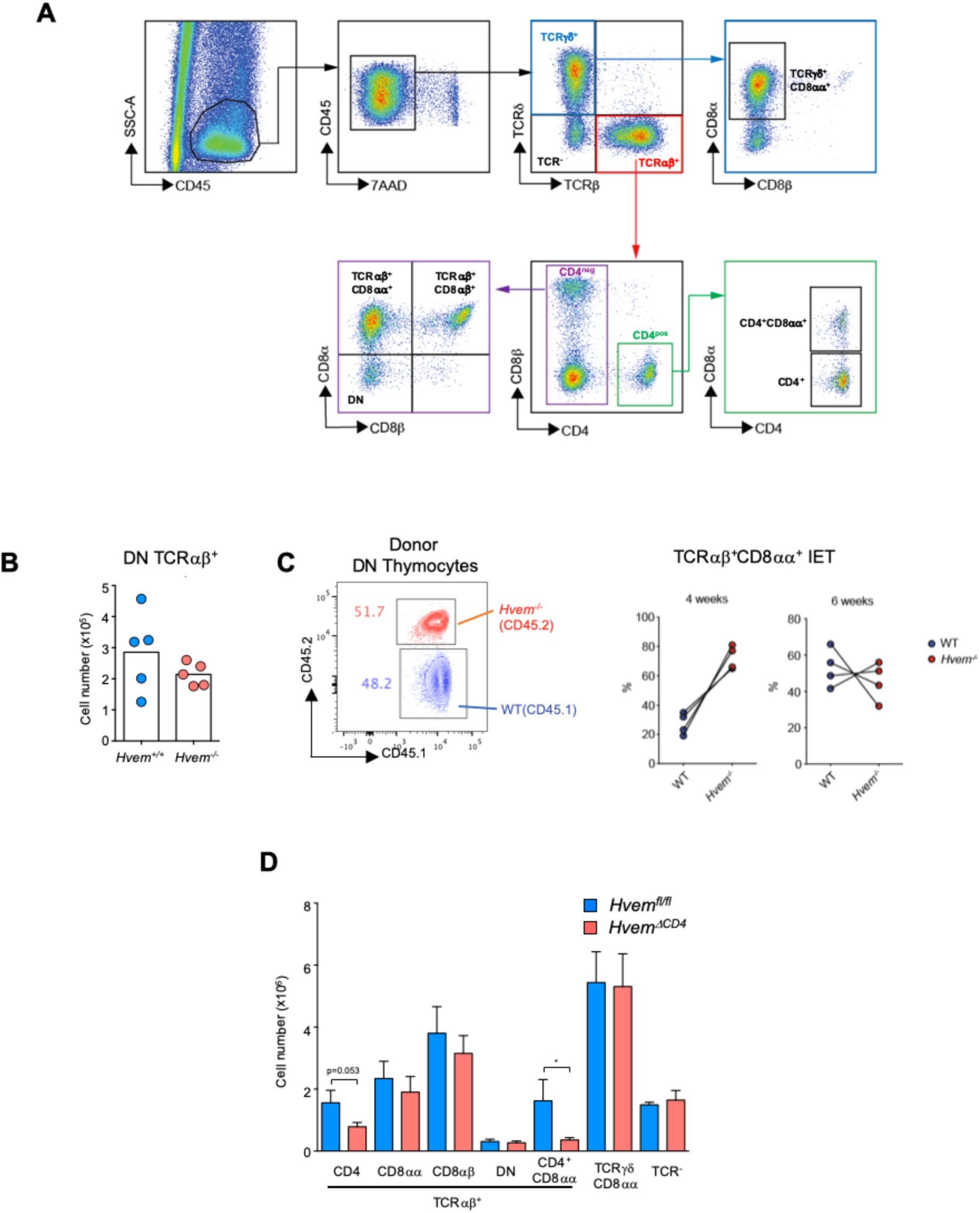
T cell intrinsic HVEM expression is not required for CD8αα IET development. (A) Gating strategy for IET subsets. Gating was performed on live, CD45^+^lymphocytes and doublets were excluded. Gating strategy for TCRγδ^+^CD8αα^+^ cells (CD45^+^TCRγδ^+^CD8α^+^CD8β^-^), for TCRαβ^+^CD8αα^+^ cells (CD45^+^TCRαβ^+^CD4^-^CD8α^+^CD8β^-^), TCRαβ^+^CD8αβ^+^ cells (CD45^+^TCRαβ^+^CD4^-^ CD8α^+^CD8β^+^), TCRαβ^+^CD4^+^CD8αα^+^ cells (CD45^+^TCRαβ^+^CD4^+^CD8α^+^CD8β^-^). (B) Absolute numbers of the precursor of CD8αα^+^ IET (CD4^-^CD8α^-^TCRβ^+^ thymocytes) in thymus from *Hvem*^*+/+*^ and *Hvem*^-/-^ mice. (C) The role of HVEM expression during CD8αα^+^ IET development in vivo. Equal numbers (1×10^5^) of sorted CD4^-^CD8^-^ NK1.1^-^ TCRβ^+^ thymocytes from CD45.1 congenic wild type and *Hvem*^-/-^ mice (CD45.2) (left) were co-transferred into *Rag1*^-/-^ recipient mice. After 4 or 6 weeks, the frequencies of CD45.1 and CD45.2 of CD8αα^+^ IET from the SI in the recipients were analyzed (right). (D) Absolute numbers of the indicated subsets in total IEL from *Hvem*^*fl/fl*^ and *Hvem*^*ΔCD4*^ mice (n=9 per each group). Statistical analysis was performed using an unpaired t-test (B, D). Statistical significance is indicated by *, p < 0.05. In B, bars show the mean and each symbol represents a measurement from a single mouse (B). Data are representative results from one of at least two independent experiments with at least four mice in each experimental group (B, C) or pooled results from at least two independent experiments with at least four mice per group in each experiment (D). Groups of co-housed littermates were analyzed.

**Figure S2. Related to Figure 2.**
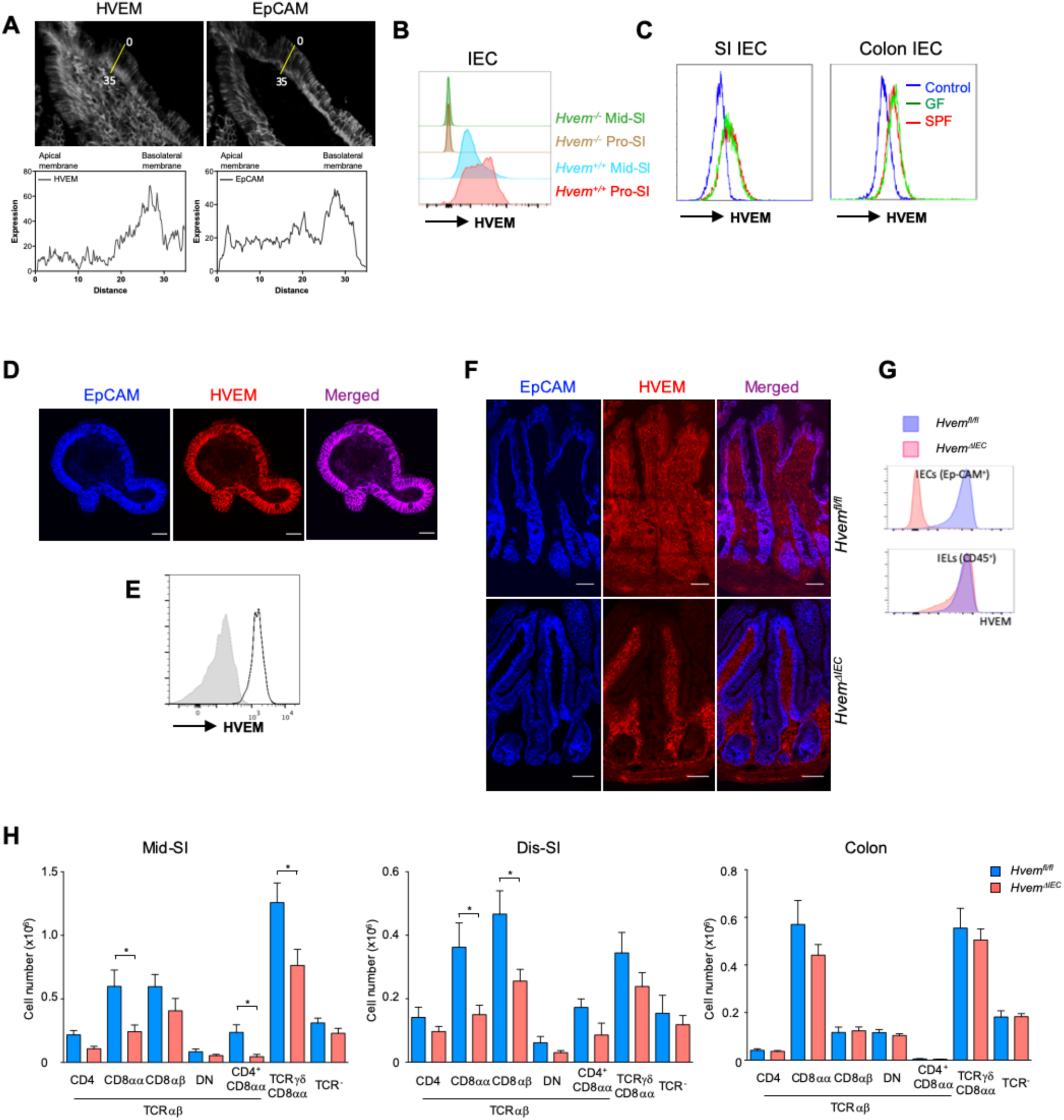
Constitutively expressed HVEM influences IET subsets. (A) HVEM is predominantly expressed on the basolateral membrane of SI IEC from wild type mice. Top: representative immunofluorescence staining of HVEM (grey) and EpCAM (grey) in the SI. Bottom: tracing of intensity of HVEM and EpCAM signal from apical to basal aspect of IEC using Fiji software. (B) HVEM expression by IEC in the middle and proximal SI of *Hvem*^*+/+*^ and *Hvem*^*-/-*^ mice. (C) HVEM expression by IEC from SI and colon from germ-free (GF) and specific-pathogen-free (SPF) mice. (D-E) Expression of HVEM in a representative organoid from proximal SI of wild type mice demonstrated with confocal microscopy (D) and flow cytometry (E). Immunofluorescence staining for HVEM (red) and EpCAM (blue) in the whole mount tissues from organoid culture. Scale bars, 50μm. (F-G) Confirmation of lack of HVEM expression on IEC in *HVEM*^*ΔIEC*^ mice using immunofluorescence staining. (F) Immunofluorescence staining for HVEM (red) and EpCAM (blue) in frozen sections of small intestine samples from *Hvem*^*fl/fl*^ and *Hvem*^*ΔIEC*^ mice. Scale bars, 50μm. (G) HVEM expression by IEC and CD45^+^ IEL from SI of *Hvem*^*fl/fl*^ and *Hvem*^*ΔIEC*^ by flow cytometry. (H) Absolute numbers of indicated subsets in total IEL from middle SI, distal SI and colon from *Hvem*^*fl/fl*^ and *Hvem*^*ΔIEC*^ mice (n=10 per group). Statistical analysis was performed using an unpaired t-test. Statistical significance is indicated by *, p < 0.05. Data represent pooled results from at least two independent experiments with at least four mice per group in each experiment. Groups of co-housed littermates were analyzed.

**Figure S3. Related to Figure 2.**
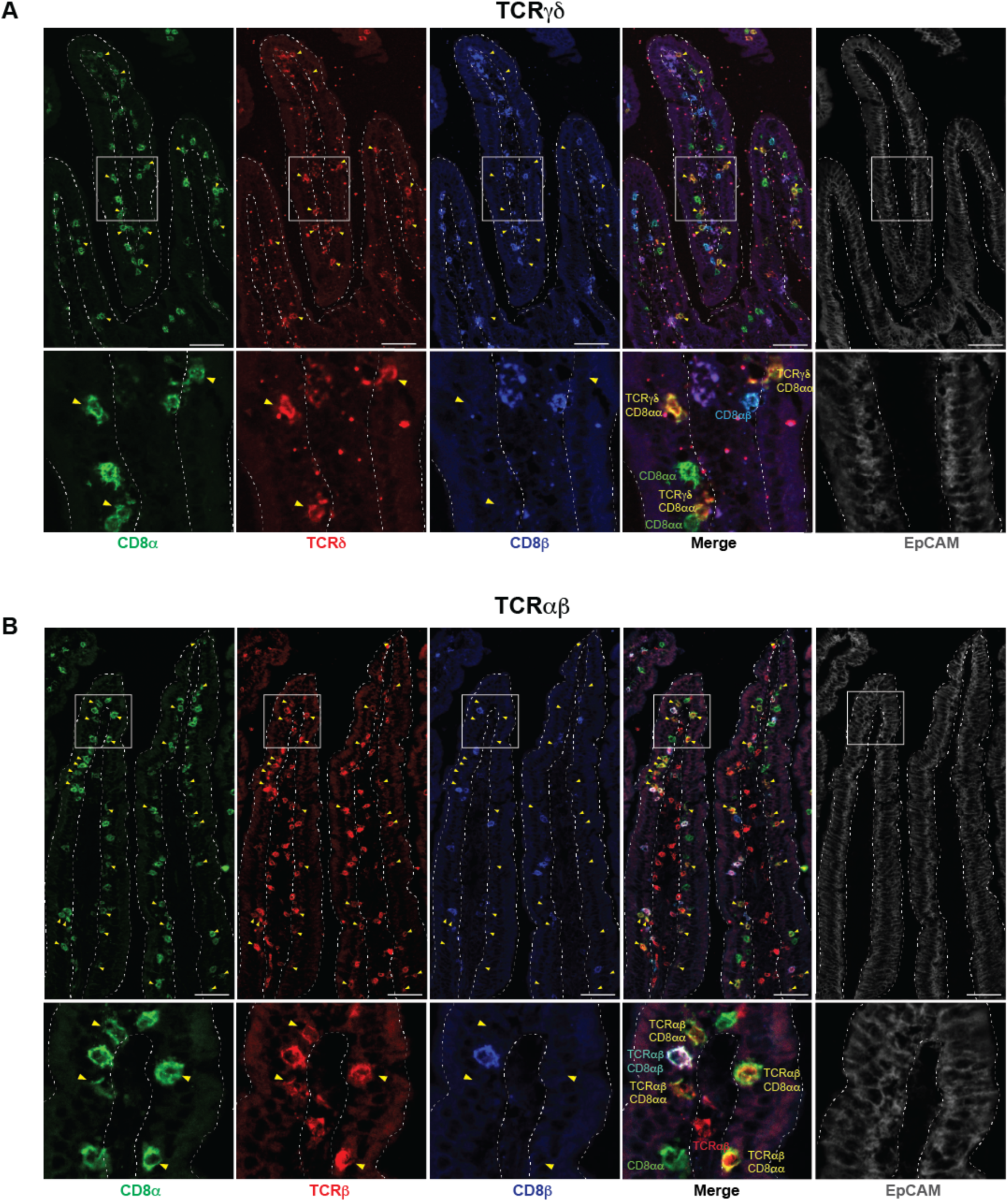
Staining strategy for immunofluorescent detection of TCRγδ IET and TCRαβ IET from proximal SI. Expanded details of areas shown in Figure 2D-2E. Immunofluorescent staining of TCRγδ IET and TCRαβ IET from proximal SI in *Hvem*^*fl/fl*^ mice. Yellow arrowheads indicate TCRγδ^+^CD8αα^+^ (A) or TCRαβ^+^CD8αα^+^ IET (B). Single channel images from proximal SI in *Hvem*^*fl/fl*^ mice detecting expression of CD8α (green), CD8β (blue), and TCRδ (red, A) or TCRβ (red, B). Expression of EpCAM shown in gray. Dashed white lines indicate the boundaries of the epithelium used for the quantitation. Scale bars, 50μm.

**Figure S4. Related to Figure 3.**
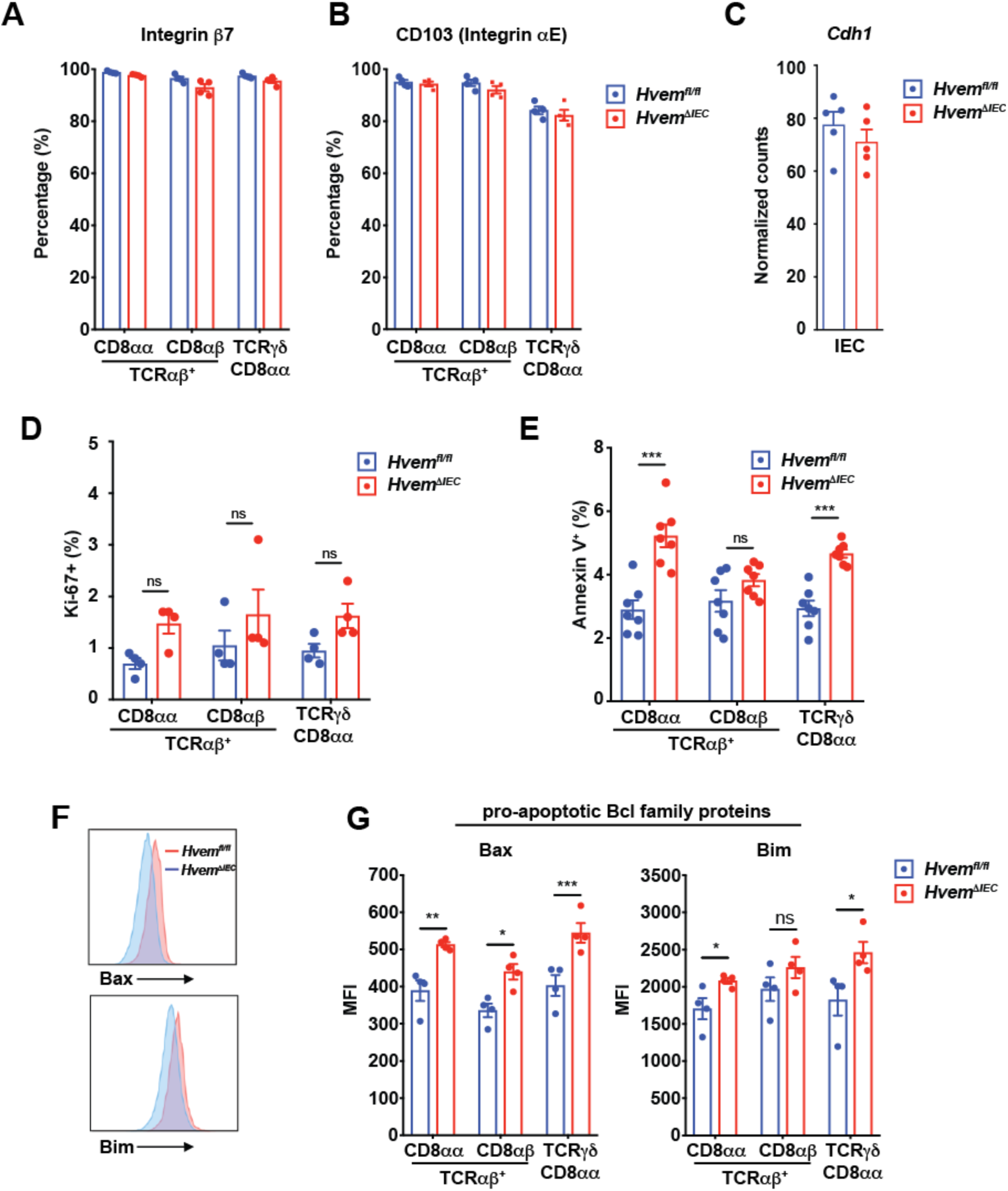
Deficiency of epithelial HVEM affects survival of CD8α^+^ IET. (A, B) Expression of Integrin β7 and αE subunits on CD8α^+^ IET subsets from proximal SI of *Hvem*^*fl/fl*^ and *Hvem*^*ΔIEC*^ mice by flow cytometry. (C) Expression of *Cdh1* mRNA by sorted IEC from proximal SI IEC of *Hvem*^*fl/fl*^ and *Hvem*^*ΔIEC*^ mice by RNA-seq. Expression levels were calculated and normalized with DESeq2. (D) Frequencies of Ki-67^+^ cells from proximal SI IET of *Hvem*^*fl/fl*^ and *Hvem*^*ΔIEC*^ mice. (E) Frequencies of Annexin V from proximal SI IET of *Hvem*^*fl/fl*^ and *Hvem*^*ΔIEC*^ mice. (F, G) Representative flow cytometry showing expression of pro-apoptotic Bcl family proteins (Bax and Bim) on TCRαβ^+^CD8αα^+^ IET from proximal SI of *Hvem*^*fl/fl*^ and *Hvem*^*ΔIEC*^ mice (F) and compiled MFI from multiple individual mice (G). Statistical analysis was performed using an unpaired t-test. Statistical significance is indicated by *, p < 0.05; **, p < 0.01; ***, p < 0.001; ns, not significant.

**Figure S5. Related to Figure 5.**
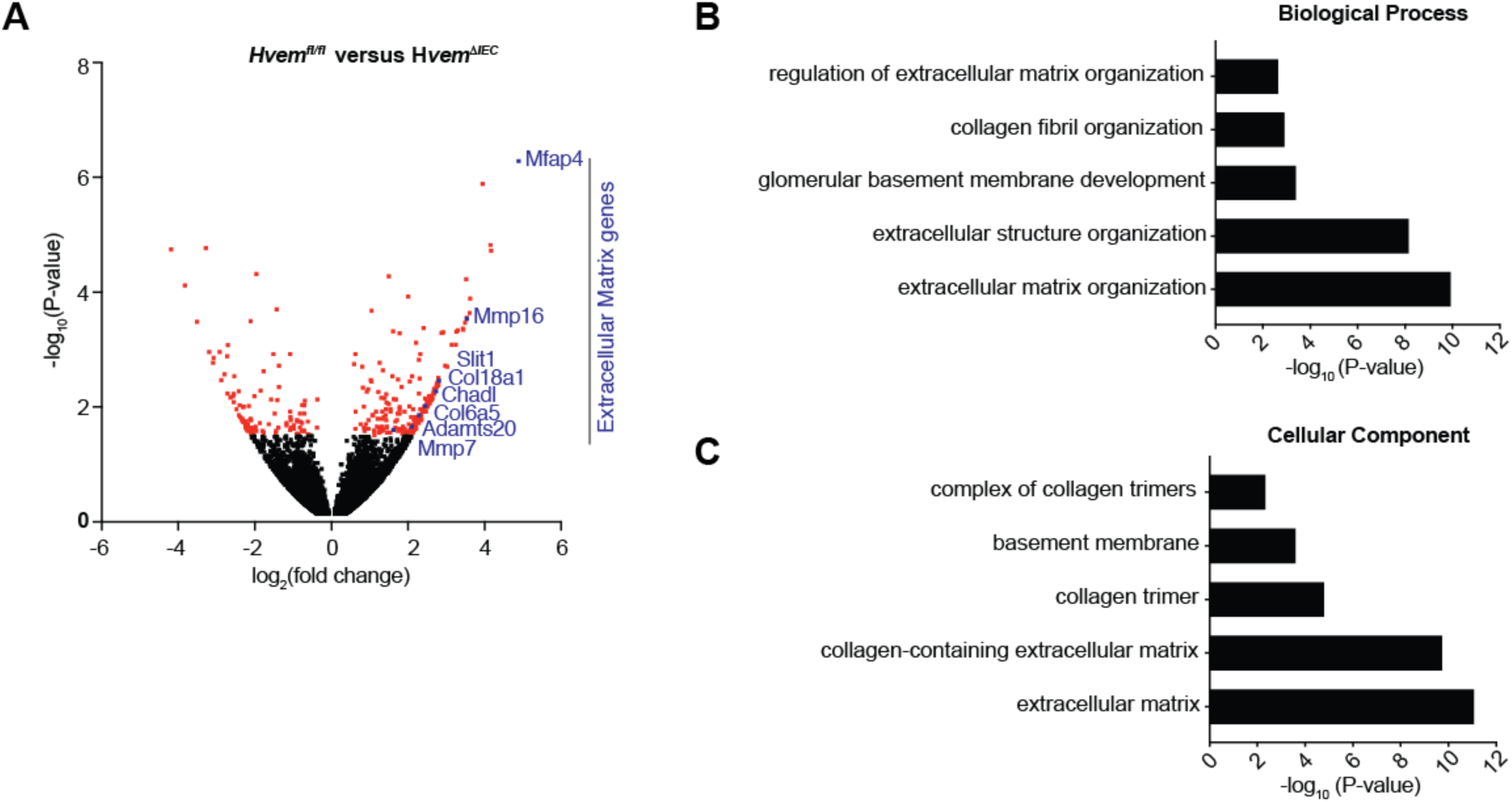
Downregulation of the genes of associated with extracellular matrix proteins in the absence of epithelial HVEM expression. (A) Volcano plots showing mean log_2_-transformed fold change (x axis) and significance (-log_10_ P value) of differentially expressed genes between the isolated IEC of the small intestine of *Hvem*^*fl/fl*^ and *Hvem*^*ΔIEC*^ mice. Extracellular matrix-related genes are indicated with blue font or blue symbol. (B-C) Isolated IEC from proximal SI of *Hvem*^*fl/fl*^ and *Hvem*^*ΔIEC*^ mice were analyzed by RNA-seq. Representative enriched GO terms related to “extracellular matrix” in the isolated IEC of the small intestine of *Hvem*^*ΔIEC*^ mice were determined by the ToppGene Suite. FDR (Benjamini-Hochberg) < 0.05 was considered statistically significant. Groups of co-housed littermates were analyzed.

