## Supplementary figures and images for "Epithelial HVEM promotes basement membrane synthesis and intraepithelial T cell survival and migration"

### graphic abstract

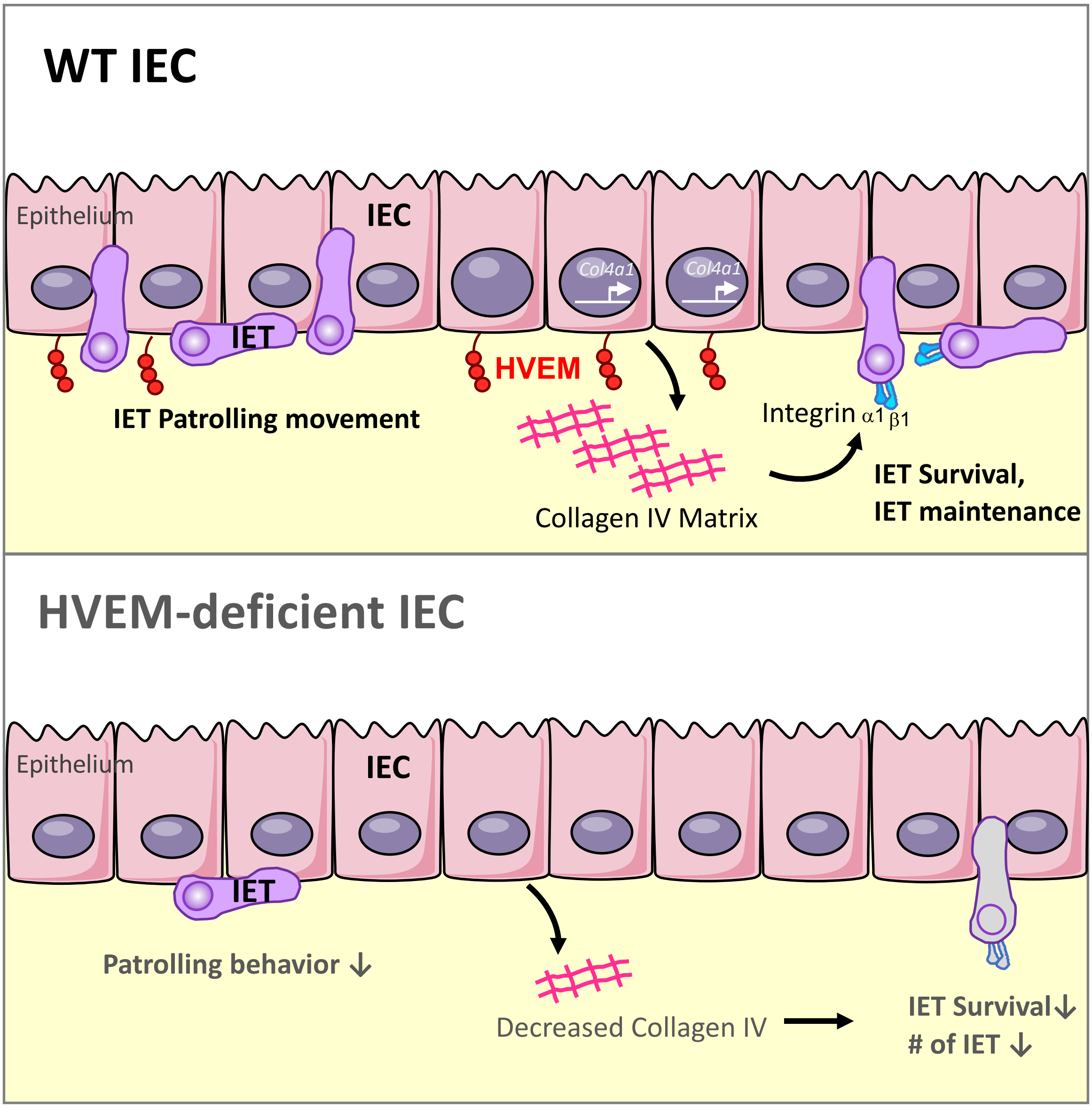
