## Supplementary material for "Epithelial HVEM promotes basement membrane synthesis and intraepithelial T cell survival and migration": highlights

**In Brief**

Intraepithelial T cells interact extensively with intestinal epithelial cells and provide for immune surveillance and barrier protection. Takahashi et al. reveal that epithelial HVEM is required for maintenance of subsets of intraepithelial T cells (IET), especially so-called natural IET that express a $\gamma\delta$ TCR or TCR $\alpha\beta$ cells that lack TCR co-receptor expression. Patrolling by IET of the epithelium also was decreased. HVEM signals to epithelium promoted synthesis of basement membrane proteins such as collagen IV. Interactions of collagen IV with integrins expressed by IET affected the survival of these cells. Therefore, these studies define a pathway induced by signals to epithelial cells that increase extracellular matrix synthesis and thereby mediate the survival and movement of resident T cells.

**Highlights**

- TNF family receptor HVEM in epithelial cells promotes survival of natural intraepithelial T cells (IET).
- Patrolling of IET is reduced when epithelial cells do not express HVEM
- Deficiency of epithelial HVEM leads to decreased epithelial synthesis of basement membrane proteins, including collagen IV.
- Collagen IV interaction with T cell $\alpha1\beta1$ integrins supports IET survival.
